# Adipose cells and tissues soften with lipid accumulation while in diabetes adipose tissue stiffens

**DOI:** 10.1101/2022.02.09.479760

**Authors:** Shada Abuhattum, Petra Kotzbeck, Raimund Schlüßler, Alexandra Harger, Angela Ariza Schellenberger, Kyoohyun Kim, Joan-Carles Escolano, Torsten Müller, Jürgen Braun, Martin Wabitsch, Matthias Tschöp, Ingolf Sack, Marko Brankatschk, Jochen Guck, Kerstin Stemmer, Anna V. Taubenberger

## Abstract

Adipose tissue expansion involves both differentiation of new precursors and size increase of mature adipocytes. While the two processes are well balanced in healthy tissues, obesity and diabetes type II are associated with abnormally enlarged adipocytes and excess lipid accumulation. We hypothesised that adipocyte shape and size changes with differentiation and lipid accumulation would be accompanied by changes in the mechanical phenotype at both the cell and tissue level. We quantified by optical diffraction tomography (ODT) that differentiating preadipocytes increased their volumes drastically. Atomic force microscopy (AFM)-indentation and -microrheology revealed that during the early phase of differentiation, human preadipocytes became more compliant and more fluid-like, concomitant with ROCK-mediated F-actin remodelling. Adipocytes that had accumulated large lipid droplets were more compliant, and further promoting lipid accumulation led to an even more compliant phenotype. In line with that, high fat diet-induced obesity was associated with more compliant adipose tissue compared to lean animals, both for drosophila fat bodies and murine gonadal adipose tissue. In contrast, adipose tissue of diabetic mice became significantly stiffer as shown not only by AFM but also magnetic resonance elastography (MRE). Altogether, we dissect relative contributions of the cytoskeleton and lipid droplets to cell and tissue mechanical changes across different functional states, such as differentiation, nutritional state and disease. Since preadipocytes are mechanosensitive, tissue stiffening in diabetes might be critical to the balance between hyperplasia and hypertrophy and moreover present a potential target in the prevention of metabolic disorders.

## Introduction

Adipose tissue is a metabolic and endocrine organ essential for whole-body homeostasis. Its most prominent cellular component, the adipocytes, are -besides other functions-specialised in the storage of energy in the form of triglycerides that can be broken down into fatty acids that provide fuel to other tissues (1). Upon changing energy status, e.g. nutrient excess or shortage, adipose tissue can be rapidly remodelled through changes in the number and/or size of adipocytes (2). Adipocyte number increase (hyperplasia) is due to adipogenesis, the differentiation of adipocyte precursor cells into mature adipocytes; while adipocyte size increase (hypertrophy) results from excess triglyceride accumulation in existing adipocytes. Both processes, hyperplasia and hypertrophy, are well-balanced in metabolically “healthy” tissue (3), while in dysfunctional tissues, such as found in obesity and diabetes type II, hypertrophic adipocytes are present in greater proportion (4)(5)(6).

Given the massive increase of the prevalence of obesity and diabetes type II over the past 50 years (7), there is great interest in the understanding of how proper adipose tissue function can be maintained during its expansion. Tissue and cell volume increase not only complicate the delivery of nutrients and oxygen, but also require excessive remodelling of the extracellular matrix (ECM) (8). Diabetes type II is often associated with tissue fibrosis (9), a state which typically manifests in altered tissue mechanical properties. Several studies have quantified the mechanical properties of adipose tissue in dependence of obesity, e.g. of mouse epididymal fat pads (10), mouse mammary fat pads (11), and human adipose tissue (12)(13). While the three mentioned studies (10)(11)(12) have not quantified mechanical changes at the cellular level, Wenderott et al. have employed AFM to probe the mechanical properties of adipose tissues from healthy and diabetic patients (13). Interestingly, they found an increase in tissue stiffness with diabetes, which was associated with enlarged adipocytes. However, only diabetic and non-diabetic obese tissues were compared, while tissues from lean and obese patients were not (13). Although mechanical alterations in diabetic tissues have been reported, it is not clear how this changed mechanical phenotype affects tissue functionality.

An altered mechanical context, as occurring with fibrosis, may also influence the differentiation of tissue-resident preadipocytes, an idea which is supported by several studies showing that adipocyte precursors are sensitive to extrinsic forces (e.g. compressive and cyclic stress (17)(16)) or mechanical cues of their microenvironment (17)(18)(19)(20). Also the mechanical properties of adipose-derived stem cells themselves have been linked to their differentiation potential, where stiffer clones showed higher osteogenic, and more compliant clones a higher adipogenic potential (21). This suggests an intrinsic relationship between the cell’s cytoskeleton and adipogenic differentiation. In fact, several studies have focused on cytoskeletal and mechanical changes in differentiating adipocytes using *in vitro* models. While undergoing adipogenesis, cells convert from an initially spread morphology towards the characteristic rounded shape of mature adipocytes that is considered a prerequisite for terminal differentiation and lipid droplet accumulation (22). These changes in cell shape and cytoskeleton have been studied recently in more detail and roles for the F-actin cytoskeleton (23)(24) and microtubules (25) have been revealed. These cytoskeletal changes can be mirrored by mechanical changes of differentiating preadipocytes, although there are controversial reports regarding mechanical alterations over the time-course of differentiation. While some studies, e.g. by Young-Nam et al. and Yu et al. showed decreasing stiffness values for differentiating preadipocytes by atomic force microscopy (AFM)(26) and micropipette aspiration (27), respectively, Labriola et al. did not detect any changes in the cells’ mechanical properties along adipogenic differentiation using AFM(28). Another study by Shoham et al. suggests that adipocyte stiffness rather increases with lipid accumulation (15). Taken together, it is incompletely understood how cytoskeletal remodelling and lipid accumulation during adipogenic differentiation affect the mechanical properties of adipose cells. Moreover, it remains unclear how surplus nutrient uptake in mature adipocytes and adipose tissue relates to cytoskeletal and mechanical changes, although this may be an important regulatory factor of cell size, differentiation of new precursors, and ultimately tissue functionality.

This putative importance of cell mechanical changes in regulating tissue functionality prompted us to investigate here the mechanical properties of adipocytes during differentiation and maturation *in vitro* and in dependence of nutritional and disease state *in vivo*. We find by AFM indentation and AFM microrheology that adipocytes become more compliant and more fluid-like with adipogenic differentiation, which is associated with a large volume increase and ROCK-mediated F-actin remodelling. Terminally differentiated cells further softened with increasing lipid droplet accumulation, which can be induced by supplementing media with monounsaturated fatty acids. In line with that, high fat diet feeding induced a more compliant mechanical phenotype in drosophila fat bodies and murine gonadal adipose tissues compared to lean counterparts as revealed by AFM. In contrast, tissue stiffness of diabetic mice was increased as revealed by AFM indentation/microrheology and magnetic resonance elastography (MRE). Taken together, our results provide new insights into the mechanical properties of adipose cells and tissue depending on differentiation and nutritional state. Since our data show that the tissue mechanical phenotype can largely differ with lipid accumulation in healthy and diabetic tissues, a better understanding of the factors causing these differences and the associated cellular responses may lead the way towards mechanisms that can be targeted to maintain the functionality of expanding fat tissues.

## Results

### Preadipocytes increase their volume with adipogenic differentiation

To monitor the adipocytes’ mechanical properties during adipogenesis, we induced human pre-adipocytes (Simpson-Golabi-Behmel Syndrome, SGBS) with an adipogenic cocktail and characterised them on day 1, 3, 6, and 11 after induction (Fig. 1A). On day 6, multiple cells with lipid droplets were seen in phase contrast and confocal images (Fig. 1B, Supplementary Fig. 1). Comparing day 6 and 11, a similar proportion of cells with lipid droplets was found (Supplementary Fig. 1), while lipid droplet sizes increased significantly on day 11 (Fig. 1C). Adipogenic differentiation and lipid droplet accumulation were accompanied by a large increase in cell volume (mean: 7,700 to 34,100 μm^3^), as quantitated by optical diffraction tomography (ODT) (Fig. 1D, E). OD tomographs also showed increased refractive indices for lipid droplets over cytoplasmic and nuclear regions as previously reported(29, 30)

**Figure 1.**
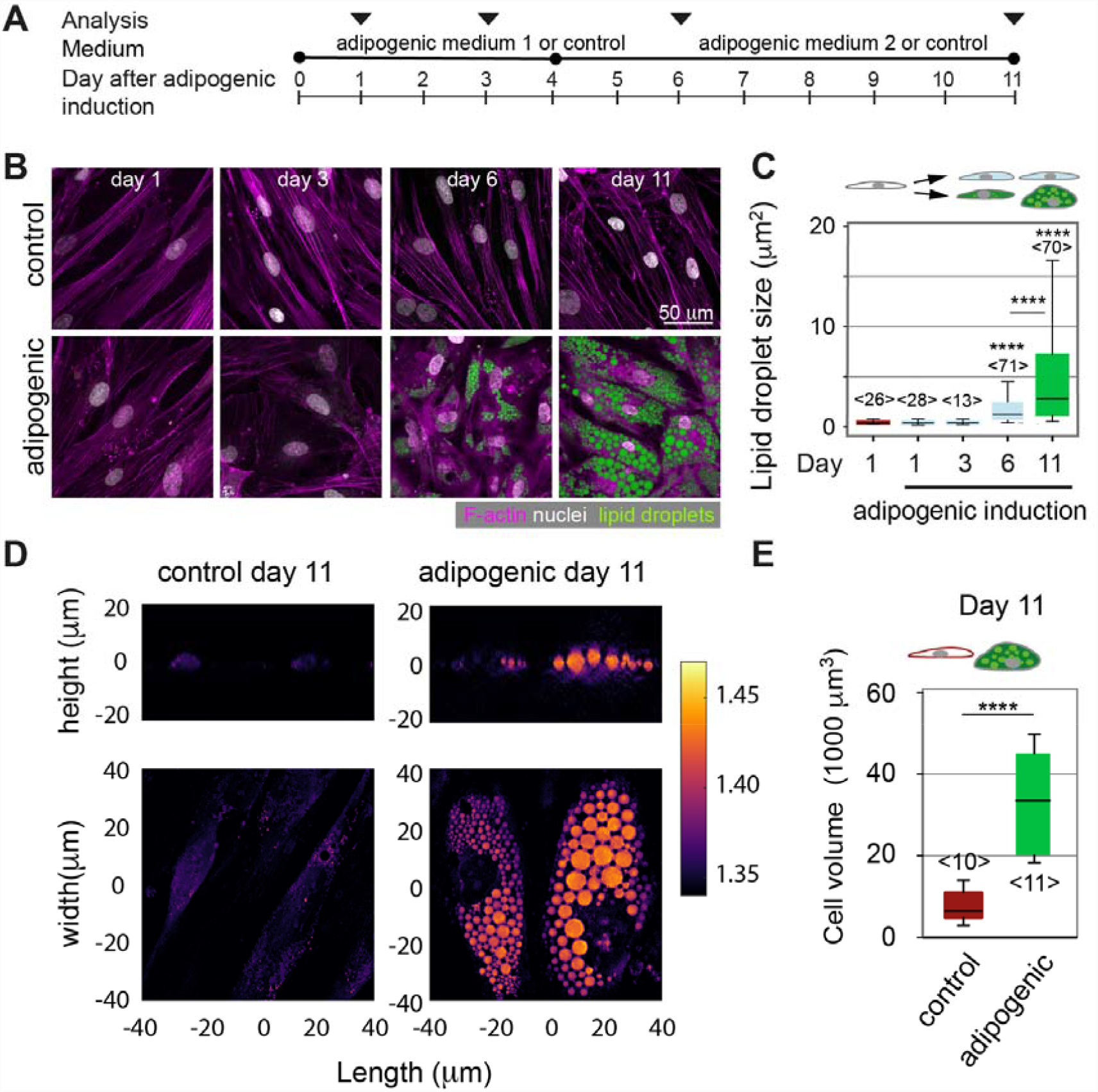
Lipid accumulation and volume changes of differentiating preadipocytes. **(A)** Schematics showing the protocol of adipogenic induction and analysis of SGBS cells. Cells were either induced with an adipogenic cocktail or kept in cell culture medium without adipogenic supplements (= control). **(B)** Representative confocal images of control cells on day 1, 3, 6 and 11 after adipogenic induction. Cells were stained for F-actin (Phalloidin-TRITC, magenta), nuclei (DAPI, white) and lipid droplets (nile red, green). **(C)** Quantification of lipid droplet size in confocal images of nile red-stained SGBS cells (cross-sectional area). The number of analysed cells is indicated in brackets <n>. Results of a statistical test (Kruskal Wallis) are shown. Comparisons to day 1 samples and specific pairs are shown. **** denotes p-values <0.0001 **(D)** Optical diffraction tomographs (ODT) of SGBS controls or adipogenically induced cells at day 11. Refractive indices are given by the shown colour scale. **(E)** Cell volumes quantified from ODT data using the Arivis software. Data are presented as box-whisker plot (25^th^, 50^th^, 75^th^ percentiles, whiskers indicate 10^th^ and 90^th^ percentiles). Number of analysed cells <n>. For comparison a Mann-Whitney test was performed. **** denotes p-values <0.0001.

### Preadipocyte stiffness decreases with adipogenic differentiation

We next set out to quantitatively assess the mechanical properties of differentiating preadipocytes by atomic force microscopy (AFM)-indentation and -microrheology (Fig. 2A). AFM-indentation revealed that within only one day of adipogenic induction, apparent Young’s moduli of induced SGBS cells significantly decreased compared to non-induced controls, indicative of more compliant cells (Fig. 2B). Apparent Young’s moduli were further reduced from day 1 to 6. Day 11 cultures, considered here as fully differentiated, displayed minimal apparent Young’s moduli on average (day 1: 3.84 ± 0.15 kPa, day 6: 0.50 ± 0.03 kPa, day 11: 0.46 ± 0.02 kPa, mean ± standard error of the mean). On both day 6 and 11, apparent Young’s moduli of cells that had accumulated lipid droplets were significantly lower compared to cells without lipid droplets (Fig. 2B). We further characterised the cells’ viscoelastic properties by AFM-microrheology, probing cells with an oscillating spherical indenter at frequencies between 3 and 200 Hz (Fig. 2C). On day 1, shear storage moduli (G’) were reduced after adipogenic differentiation compared to respective non-induced controls, while shear loss moduli G’’ remained comparable. This shows a reduced elastic resistance of the cells to deformation after their adipogenic induction, which is in line with the above-described indentation results showing a decrease in apparent Youngs moduli after adipogenic induction. On day 11, both shear storage and loss moduli of adipogenically induced cells were drastically reduced compared to non-induced cells. At low frequencies below 10Hz, the observed changes in shear storage and loss moduli of adipogenically induced cells resulted in higher mean loss tangents (defined as G’’/G’) (Fig. 2C) compared to non-adipogenic controls (both day 1 and 11). The increased loss tangents are indicative of a more fluid-like response to cell deformation upon adipogenic differentiation. Loss tangents obtained for the different conditions changed relatively towards higher frequencies (e.g. 200Hz), with day 11 adipogenic showing the lowest levels and thus a more elastic response. Similar to SGBS cells, apparent Young’s moduli of NIH3T3-L1 cells were similarly decreased after adipogenic differentiation (Supplementary Fig. 2).

**Figure 2.**
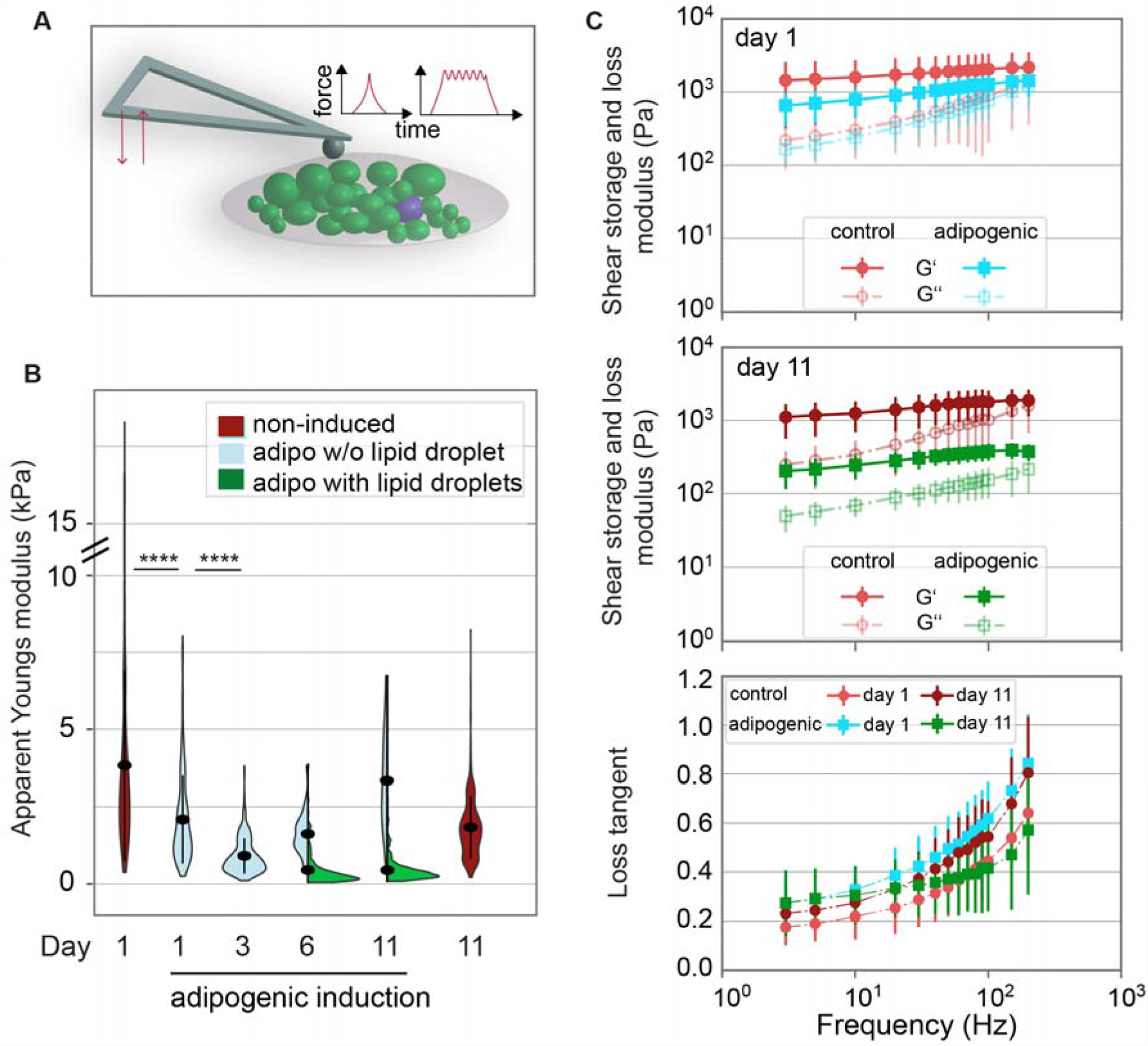
Mechanical characterisation of differentiating preadipocytes. **(A)** Schematics showing the mechanical characterisation of adipocytes using AFM indentation and microrheology measurements. For indentation tests, the indenter (bead of 5μm diameter) was lowered onto a single cell using a piezo extension speed of 5μm/sec until a preset force setpoint of 2nN was reached. From the indentation part of the force-distance curve, the apparent Youngs modulus was extracted. For microrheology, after reaching the force setpoint, the piezo elements were oscillated using a sinusoidal signal at defined frequency between 3 and 300Hz and at an amplitude of 10nm. Analysis of the force/indentation amplitudes and phase angle yielded G’ and G’’. **(B)** Mechanical characterisation of SGBS cells by AFM indentation experiments. Apparent Young’s modulus data are presented as violin plots (generated in R). From day 6 of adipogenic differentiation, cells with lipid droplets (green) or without lipid droplets (light blue), according to visual inspection of phase contrast images, are distinguished and presented in asymmetric violin plots (blue, green). For each timepoint and media condition, at least 390 cells were probed. For comparison a Kruskal-Wallis test with a Dunn’s multiple comparison test was performed (whole datasets for each day). **** denotes p-values <0.0001 (only tests between selected pairs are shown). **(C)** AFM microrheology results on adipogenically induced (turquoise, green) and control SGBS cells (red, rose) on day 1 (top) and 11 (middle). Mean ± SEM are shown, filled symbols correspond to shear storage moduli G’, unfilled symbols to loss moduli G’’. Below: As a measure of fluidity, loss tangents (calculated as G’’/G’) are plotted over frequency.

### Decreased ROCK activity promotes adipogenic differentiation

Since we found that the apparent Young’s moduli decreased already before evident accumulation of lipid droplets, we hypothesized that the cell cytoskeleton was directly modulated as an early effect of adipogenic induction. We therefore characterised the F-actin cytoskeleton in SGBS cells at different timepoints after adipogenic induction. While cells initially showed prominent F-actin stress fibrils, these were reduced in number and staining intensity during adipogenic differentiation (Fig. 3A,B). The decreased anisotropy coefficient indicated a more random F-actin fibril orientation starting from day 1 of adipogenic differentiation (Fig. 3C), which is in line with previous reports showing a role of F-actin modulation on adipogenic differentiation (23). To test the effect of F-actin on cell stiffness at different timepoints of differentiation, we next treated differentiating preadipocytes with the Rho-associated kinase (ROCK) inhibitor Y27632 and F-actin depolymerising drug cytochalasin D (Fig. 3D,E). In non-adipogenic controls, both ROCK and F-actin inhibition led to significantly reduced apparent Youngs moduli. In contrast, after adipogenic induction this inhibitory effect was blunted. In addition, already at day 1 of adipogenic differentiation, quantitative Western blot analysis indicated a decrease in myosin light chain mono (pMLC: Ser19) - and di-phosphorylation (ppMLC: Thr18/Ser19 (Fig. 3F, Supplementary Fig. 3), which is in line with the previously seen drop in cell stiffness, since myosin-2 motors generate contractile tension in the F-actin network (31). Thus, the apparent Young’s modulus decrease with adipocyte differentiation might be due to ROCK-dependent modulation of the F-actin network and/or contractility. RhoA and its downstream target ROCK have been associated with commitment of mesenchymal stromal cell differentiation towards the adipogenic lineage (24)(23). We therefore treated differentiating SGBS cells with Y-27632 and cytochalasin D from day 1 of adipogenic induction and assessed the effect on cell differentiation on day 6 and 11 by comparing oil red stainings of controls or inhibitor-treated cultures (Fig. 3G, H). A higher degree of adipogenic differentiation with drug treatment, in particular Y-27632, was found, which was supported by the larger number of lipid droplet containing cells (Fig. 3I, Supplementary Fig. 4). We also found increased levels of PPAR*γ*, a master regulator of adipogenesis in Western blot lysates after ROCK inhibitor treatment (Supplementary Fig. 4). Altogether, these results underline the crucial role of F-actin remodelling and as a consequence the cells’ mechanical phenotype in adipogenesis, in particular at its early stage.

**Figure 3.**
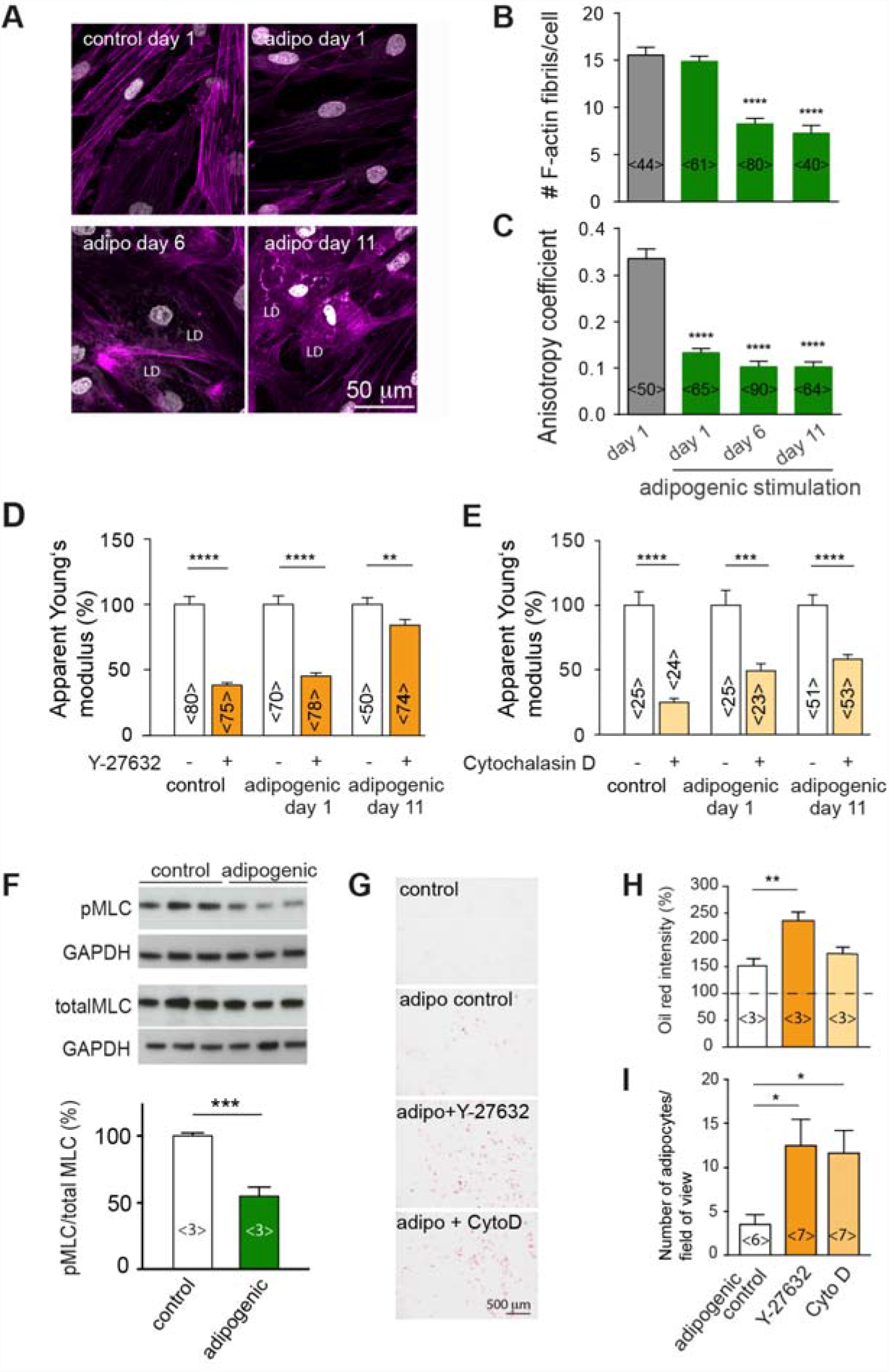
F-actin cytoskeleton changes during adipogenic differentiation. **(A)** F-actin stress fibrils at different time points, visualised by Phalloidin-TRITC (magenta). **(B)** Quantification of the number of stress fibrils per cell in confocal microscopy images using Fiji at different timepoints after adipogenic differentiation. **(C)** Anisotropy coefficient determined using Image J. Data in **B** and **C** are presented as mean ± SEM. Number of analysed cells are indicated in brackets <n>. Datasets were compared using a one-way ANOVA test combined with a Dunnett’s multiple comparison test conducted in Graphpad Prism. **** denotes p-values <0.0001. **(D + E)** Apparent Young’s moduli of SGBS cells treated with 10 μM ROCK inhibitor Y-27632 or 400 nM F-actin depolymerising drug cytochalasin D (Cyto D) at day 1 and 11 of adipogenic differentiation and non-adipogenic controls. Apparent Young’s moduli were normalised to respective untreated controls. Mean ± SEM are presented. Datasets were compared to non-adipogenic controls using a Mann-Whitney test conducted in Graphpad Prism. ** denotes p-values <0.01, *** denotes p-values <0.001, **** denotes p-values <0.0001. **(F)** Levels of phosphorylated myosin light chain (pMLC: Ser19) relative to total myosin of controls and adipogenically induced cells (day 1). pMLC and ppMLC levels were normalised to total MLC and GAPDH loading controls after densitometric analysis in Fiji. Uncropped blots are shown in Supplementary Fig. 3. **(G)** Brightfield images of oil red-stained SGBS cells differentiated in absence (control) or presence of 10 μM ROCK inhibitor Y-27632 or 400 nM cytochalasin D for 6 days. **(H)** For quantitative comparison, oil red was extracted and the absorption signal read. The absorption value was normalised to non-adipogenic controls (dotted line). Mean ± SEM of three independent experiments are presented and results of a t-test are indicated. ** denotes p-values <0.01. **(I)** Quantification of cells with lipid droplets using confocal images of nile red and DAPI stained SGBS cells 11 days after adipogenic differentiation. Mean ± SEM are presented and results of a t-test comparing controls to Y-27632 or cytochalasin D treated samples. The number of analysed samples is given by <n>. * denotes p-values <0.05.

**Figure 4.**
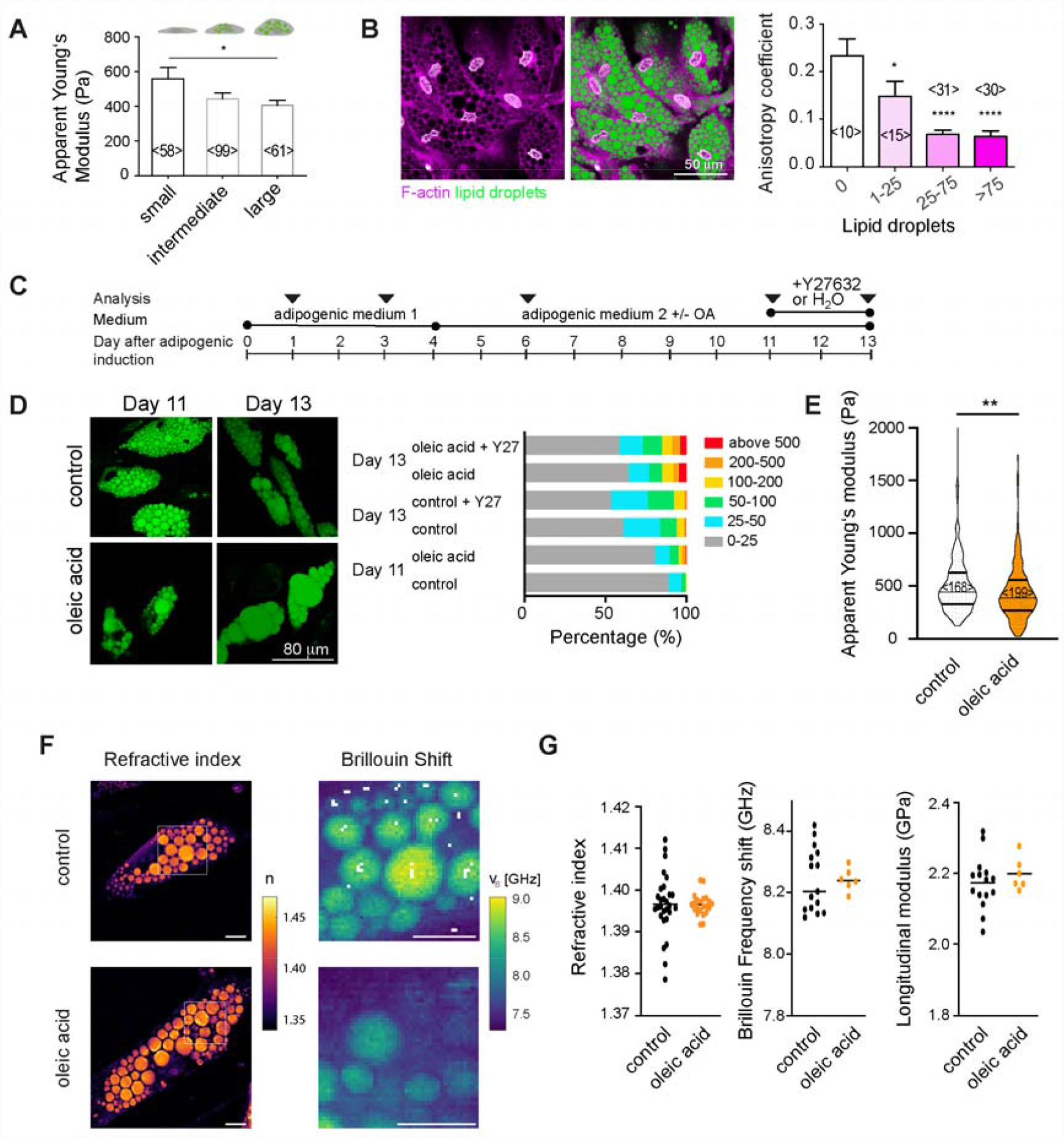
Cell mechanical changes during feeding. **(A)** Apparent Young’s moduli for cells with small, intermediate and large lipid droplets. Mean ± SEM is shown. **(B)** Confocal microscopy image of SGBS cells at day 11 after adipogenic induction (F-actin: Phalloidin-TRITC, magenta; nuclei: DAPI, white; lipid droplets: nile red (green). Anisotropy coefficient for cells depending on lipid droplet sizes (day 11 of adipogenic). The numbers of analysed cells are indicated <n> **(C)** Schematics showing culture conditions for SGBS cells. **(D)** Nile red stained SGBS cells on day 11 and 13 after oleic acid feeding (60 μM from day 4 on). Controls were fed with equal concentrations of the carrier BSA (fatty acid free). Right: Classification of lipid droplet sizes (in μm^2^) (analysed using Ilastik software) **(E)** Violin plots showing apparent Young’s moduli of control or oleic acid/treated cells (day 11). Data were compared by a Mann-Whitney test <n> represents the number of probed cells. **(F)** Representative ODT images (refractive index n) and Brillouin maps of selected regions (Brillouin shift v_B_) of controls and OA treated cells (day 11). Scale bar 10 μm. **(G)** Scatter plots showing refractive indices, Brillouin frequency shifts and longitudinal moduli of lipid droplets. Individual dots represent average values of one analysed cell. Lines indicate averages of different cells. **** denote p-values <0.0001, ** p-values<0.01, * p-values<0.05.

### Adipocytes with enlarged lipid droplets are less stiff in vitro

Having observed changes of the cells’ mechanical phenotype, as associated with cytoskeletal changes, with differentiation, we aimed to elucidate whether also excessive lipid uptake and associated lipid droplet size increase would lead to changes in the mechanical properties of mature preadipocytes. Thus, we characterised the cell mechanical properties of mature adipocytes depending on lipid droplet accumulation. On day 11 after adipogenic induction, cells with enlarged lipid droplets showed decreased apparent Young’s moduli compared to cells with small lipid droplets (Fig. 4A). In addition, F-actin was found to be more randomly distributed in cells with more lipid droplets (Fig. 4B). To further compare the effect of lipid droplet accumulation on cell mechanics, we supplemented SGBS cell medium with oleic acid (OA) (Fig. 4C), which resulted in enlarged lipid droplets (Fig. 4D). Treating cells during OA feeding with the ROCK inhibitor Y27632 (10 μM) had no significant effect on lipid droplet size distribution (Fig. 4D, Supplementary Fig. 5). OA-fed adipocytes had significantly reduced apparent Young’s moduli as revealed by AFM-indentation tests (Fig. 4E). To obtain insights into the optical and mechanical properties of intracellular compartments including lipid droplets, we analysed mature adipocytes (control and OA fed) by a combined epifluorescence, ODT and Brillouin microscopy (FOB) setup (29)(Fig. 4F,G). FOB revealed clearly increased values for refractive index and Brillouin frequency shift for lipid droplets compared to cytoplasmic regions (Fig. 4F), which relates to decreased longitudinal moduli for lipid droplets as also recently shown quantitatively (29). No significant differences of refractive indices and Brillouin frequency shifts were found though when comparing the lipid droplets of control and OA-fed adipocytes. Calculating the longitudinal moduli of lipid droplets, no significant differences between controls and OA fed adipocytes were seen (Fig. 4G). This suggests that mechanical changes upon OA feeding revealed by AFM indentation tests are either explained by a relatively increased volume of the more compliant lipid droplets (compared to the cytoplasm), or are dominated by other cellular components than lipid droplets, e.g. the cell cortex.

### Adipose tissue softens after high fat diet-induced obesity

Next, we were interested to see whether the observed cell mechanical changes upon “feeding” cells *in vitro* were also translated to the tissue level. Two different animal models were used: fat bodies from drosophila larvae and gonadal adipose tissue from C57BL76J mice. Fat bodies from larvae fed with lipid-free basal diet supplemented with olive oil (=high fat diet, HFD) showed enlarged lipid droplets with respect to controls kept on basal diet supplemented with ß-Sitosterol (=controls, Fig. 5A,B). In addition, we found that apparent Young’s moduli detected for the fat bodies of HFD larvae were significantly decreased with respect to their control siblings (=chow, Fig. 5C). We also compared the mechanical properties of lean and obese gonadal white adipose tissue (GWAT) from wildtype C57BL76J mice that had been fed a normal (chow) or HFD (Fig. 5D). GWAT of HFD-fed mice comprised adipocytes of increased size when compared to tissue from chow-fed mice (Fig. 5E). AFM-indentation tests revealed lower apparent Young’s moduli for HFD-compared to chow-fed mice (Fig. 5F). When dynamically comparing the mechanical phenotype of HFD and chow-fed mice by AFM microrheology, depending on the frequency, decreased shear storage but similar shear loss moduli were obtained for HFD compared to chow-fed mice. Moduli showed a distinct frequency-dependence (Supplementary Fig. 6). At whole tissue level and with higher frequencies, magnetic resonance elastography (MRE) revealed similar shear storage and loss moduli for HFD and chow-fed mice (Supplementary Fig. 6). Together, these results suggest that at the single cell level adipose tissue rather decreases its stiffness with excessive accumulation of lipids induced by HFD, which is in line with the above-described *in vitro* feeding experiments but appears at first sight contrary to previous reports on obese (and diabetic) tissue samples (13). Thus, we next wished to analyse not only adipose tissues from healthy obese but also diabetic mice.

**Figure 5.**
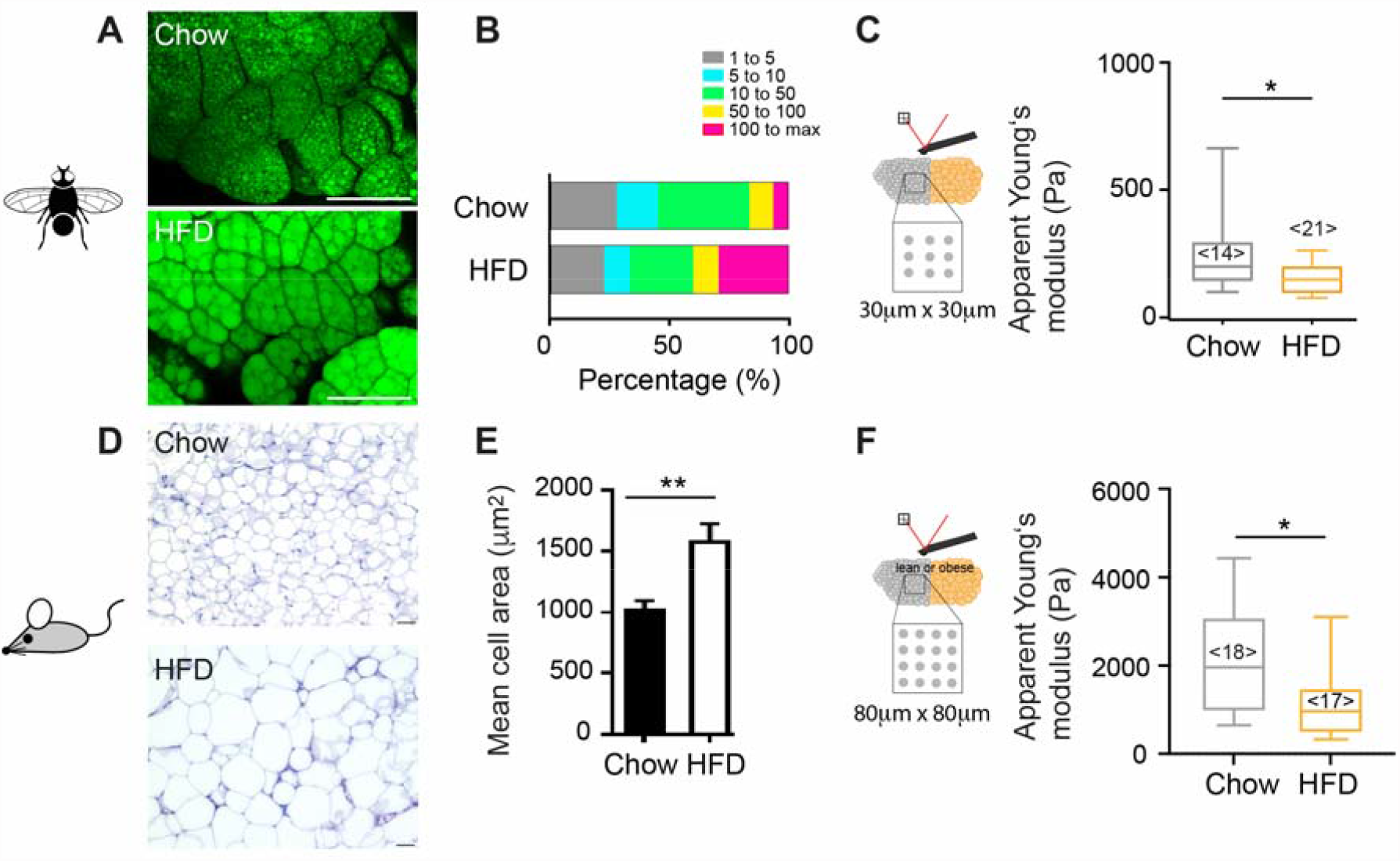
Tissue mechanical changes during feeding. **(A)** Confocal microscopy images of drosophila fat bodies of animals fed on control (top) and lipid-rich (below) diet (lipid droplets stained with bodipy-505, green). Scale bar 100 μm **(B)** Quantification of lipid droplet sizes (in μm^2^) of drosophila fat bodies (n= 3 animals) in Fiji. **(C)** Boxplots showing apparent Young’s moduli of fat bodies, obtained by AFM indentation tests using a spherical indenter of 5μm radius on multiple positions. Data were compared by a Mann Whitney test. <n> represents the number of probed tissues. **(D)** H&E stainings of gonadal adipose tissue of lean and obese mice (chow and high fat diet HFD). **(E)** Quantification of cell sizes from histology stainings (in μm^2^). **(F)** Boxplots showing apparent Young’s moduli of gonadal adipose tissue, obtained by AFM indentation tests using a spherical indenter of 5μm radius on multiple positions. Datasets were compared using a Mann-Whitney test. ** denotes p-values <0.01, * p-values <0.05.

### Adipose tissue stiffness increases in diabetic mice

We then characterised the mechanical properties of the adipose tissue in db/db mice. This mouse model carries a loss of function mutation in the leptin receptor gene and provides a well-established model of obesity-induced type 2 diabetes. These db/db mice typically develop first signs of hyperglycemia at an age of 8-10 weeks (32). Lean counterparts were control mice of the same dock 7 background as db/db mice but with wildtype leptin receptor (ctr). Paraffine sections of the gonadal adipose tissue of 4 months old animals showed that the adipocyte size in db/db mice was enlarged compared to the WT controls (Fig. 6A) and stained positive for the macrophage marker CD68, indicative for adipose tissue inflammation (Fig. 6B). On average, db/db mice also were characterised by elevated blood glucose levels, unlike control dock 7 mice, and lean and HFD-fed wildtype mice (Supplemental Fig. 7). Unlike the diet-induced obese mice, increased apparent Young’s moduli were detected for obese diabetic (db/db) mice compared to lean controls (Fig. 6C). In contrast, adipose tissue of another genetically obese but non-diabetic mouse strain (ob/ob, leptin-deficient) was comparable to respective lean controls (Supplementary Fig. 7). Moreover, a higher variance of apparent Young’s moduli of db/db mice was observed compared to lean control animals, indicating a higher level of heterogeneity of local tissue stiffness (Fig. 6C). To characterise the viscoelastic properties of the adipose tissue, AFM-microrheology and magnetic resonance elastography (MRE) measurements were employed, probing tissue stiffness at a local and a more global scale, respectively. Both techniques, albeit operating in different frequency ranges, revealed increased shear storage moduli (G’) for db/db mice compared to controls (Fig. 6D,E). Thus, in conclusion, the seen mechanical changes at the single cell level, i.e. softening with lipid accumulation, were only partly seen in high fat diet induced obese (healthy) tissues but not in tissues of diabetic animals. This suggests that tissue mechanical properties are unlikely to be only explained by the mechanical properties of constituent adipocytes, but may be influenced by additional factors, e.g. increased matrix deposition.

**Figure 6.**
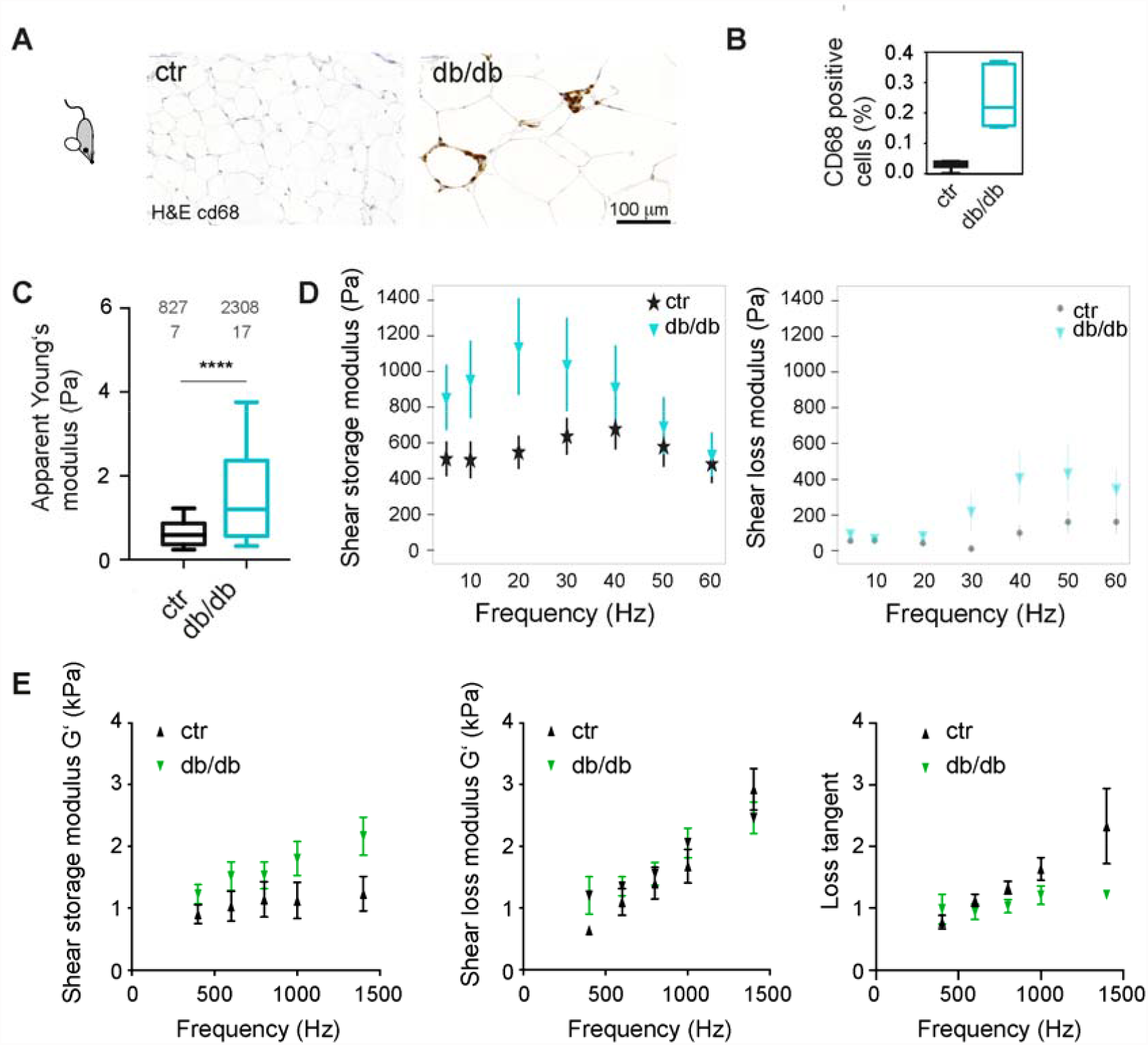
Tissue mechanical changes in diabetic mice. **(A)** H&E and immunostainings for CD68 of gonadal adipose tissue of lean control and obese mice (dock 7 (top) and db/db (below) mouse strain). **(B)** Quantification of CD68 positive cells. **(C)** Boxplots showing apparent Young’s moduli of gonadal adipose tissue, obtained by AFM indentation tests using a spherical indenter of 5μm radius on multiple positions. Datasets were compared using a Mann-Whitney test. **** denote p-values <0.001. **(D)** AFM microrheology on gonadal adipose tissue of lean (dock7, n=5) and obese (db/db, n=3) mice over a frequency rate of 5 to 60 Hz. Mean +/-standard deviation are shown **(E)** Magnetic resonance elastography on gonadal adipose tissue of lean (dock7, n=6) and obese, diabetic (db/db, n=6) mice over a frequency rate of 400 to 1400 Hz. Mean +/-standard error on the mean is shown.

## Discussion

### Preadipocyte stiffness decreases during adipogenic differentiation

We have investigated the mechanical changes of preadipocytes during the time-course of differentiation and in response to nutrient uptake. In addition, we studied differences in the mechanical properties of the gonadal adipose tissue from lean, diet-induced obese, and diabetic db/db mice. *In vitro* and *in vivo*, the increase of cell and lipid droplet sizes was associated with decreased stiffness. In detail, we first showed decreasing stiffness of differentiating adipocytes, which is associated with changes in the F-actin cytoskeleton (Fig. 2). Within only 1 day after stimulation with an adipogenic cocktail, we detected a drop in cortical stiffness of SGBS cells, which we attribute to the action of the adipogenic cocktail. When we systematically tested the effect of single components of the adipogenic cocktail on cell stiffness, we found a similar drop in cell stiffness upon treatment with IBMX as well as morphological changes in the F-actin network resembling the one of adipogenically induced cultures (Supplementary Fig. 8). IBMX elevates cAMP signalling and can thereby accelerate initiation of the adipogenic differentiation program (33). This effect of cAMP signalling was recently attributed to the activity of the cAMP-dependent protein kinase A (PKA) causing downregulation of Rho A (34). Rho/ROCK are known to affect actin cytoskeleton remodeling (35)(36) and actomyosin contractility (37)(38). In principle, both processes could explain the decreased cell stiffness we observed. Under the action of the adipogenic cocktail we indeed detected decreased levels of pMLC. In addition, probing differentiating cells under the effect of F-actin cytoskeleton perturbing drugs suggested that the F-actin cortex became less stiff at the later timepoints, e.g. due to a less ordered or less dense F-actin network (39). In line with this, Labriola et al. recently reported a less dense F-actin network for adipogenically differentiated, adipose tissue-derived stem cells (28). Treating cells with a ROCK inhibitor in addition to adipogenic differentiation further promoted adipogenic differentiation. This is consistent with previous studies that have shown that F-actin disassembly by expression of dominant negative RhoA (24)(23) or cytoskeletal drugs promotes adipogenic differentiation in human mesenchymal stem cells (23) and mouse embryonic stem cells (40) and fibroblasts (41). Konubi et al. have investigated the transition of stress fibril breakdown towards cortical F-actin structures in more detail and identified a role for insulin-PI3K-Rac1 signaling in stabilising cortical actin structures that are required for the completion of adipogenic differentiation. As possible link between cytoskeleton regulation and expression of adipogenic genes, transcriptional coactivators such as YAP1(42) or megakaryoblastic leukemia 1 (MLK1) (23) have been suggested. Together, it is interesting and important that pathways driving adipogenic differentiation can be modulated by cell mechanical/cytoskeletal cues.

### Preadipocytes decrease their stiffness with lipid accumulation

Over the time-course studied (up to 13 days), mature adipocytes with large lipid droplets were characterised by low apparent Young’s moduli. At day 6 and 11, two cell populations could be mechanically and morphologically distinguished: differentiated adipocytes with lipid droplets were characterised by lower stiffness, while cells that had not differentiated (as indicated by absence of lipid droplets) were significantly stiffer. Previously, there have been a handful of studies on the mechanical properties of differentiating murine preadipocytes (30)(26), and mesenchymal stromal cells (21)(28), with partly controversial results. Using AFM, Young-Nam et al. detected a decrease in cell stiffness in differentiating NIH-3T3 cells, which is consistent with our results (26). Gonzalez-Cruz et al., using AFM stress relaxation tests on rounded cells, reported that within a heterogenous population of adipose-derived stem cells, the more compliant clones had a higher potential for adipogenic differentiation (21). In contrast, stiffer clones had a higher potential to differentiate towards the osteogenic lineage, underlining that the cellular mechanical phenotype was predictive of adipose-derived stem cell differentiation. In another study (28), the authors followed the mechanical properties of adipogenically induced and control cells over a time-course of 14 days, and used molecular beacons to correlate PPAR*γ* expression to mechanical phenotype. While PPAR*γ* expression was associated with a more compliant phenotype compared to PPAR*γ* negative cells, there was no change in stiffness with time when only PPAR*γ* positive cells were analysed. Therefore, they suggested that over time there is an accumulation of the intrinsically more compliant PPAR*γ* positive cells, but single cells, once committed to adipogenic differentiation, would not change their mechanical phenotype. We also detected initially a high variability in apparent Young’s moduli for undifferentiated cells, which may reflect differences in adipogenic differentiation potential within the whole population. The stiffness drop after 1 day of adipogenic induction is, as discussed above, possibly attributed to Rho/ROCK inhibition through the adipogenic cocktail. With differentiation, we detected significant differences between a pool of cells that did not differentiate and differentiated cells that accumulated lipid droplets. Our finding that the fraction of non-differentiated cells became stiffer from day 3 to 11 may be attributed to the influence of IBMX. In addition, at earlier timepoints (days 3 and 6) there might be cells on their way to differentiation that are compliant but have not accumulated lipid droplets yet. This fraction is expected to become smaller, once more and more cells differentiate, leaving behind stiffer cells that may have only a low or no potential for adipogenic differentiation. Thus, our results of cells becoming gradually more compliant with time could be partly explained by an accumulation of adipogenically committed cells in the pool of analysed cells. However, analysing the distribution of day 1 controls and day 6/11 samples in more detail, there is a clear shift of the whole population and hardly no overlap between the distributions (Fig. 2B). From this we conclude that the observed increased deformability of cells with time after adipogenic induction, and in particular with lipid droplet accumulation, is not simply an enrichment of an early existing compliant population; there appears to be an additional decrease in stiffness with lipid droplet accumulation. This is further supported by the finding of larger lipid droplets correlating with more compliant cells, as seen on day 13 (Fig. 3C-E). What is the relative contribution of the lipid droplets, accumulating during maturation, to cell mechanics? Shoham et al. conducted AFM-indentation tests on differentiated NIH3T3-L1 cells and detected increased stiffness values when probing the cells over the nucleus rather than over lipid droplets (30). Using a finite element modelling approach, they conclude that accumulation of lipid droplets, presumably stiffer compared to cytoplasm, would lead to increased stiffness of mature adipocytes. However, no direct measurements in dependence of lipid droplet size and lipid droplet accumulation over time were performed, which makes these results difficult to compare directly with our study. We suppose that in our AFM-indentation tests (indentation depths <2 μm) we predominantly probed the cell cortex, and less likely the underlying lipid droplets. Thus, our data on oleic acid-fed adipocytes showing decreased apparent Young’s moduli for cells with enlarged lipid droplets likely reflect changes in the F-actin cortex of the cells. This is in line with a previous study that has compared the F-actin levels in the cortex of oleic acid fed adipocytes (43). This study has reported a 50% decrease in F-actin cortical levels for oleic acid fed cells that -similar to our experiments-showed enlarged lipid droplets. Interestingly, decreased cortical F-actin was shown in that study to be associated with lower glucose uptake (43). It is well acknowledged that GLUT4 translocation to the plasma membrane (required for insulin dependent glucose uptake) is dependent on cortical F-actin (44). Thus, lipid droplet configuration (multiple versus unilocular), insulin sensitivity and F-actin cytoskeleton appear to be tightly coupled. Mechanical probing of mature adipocytes may therefore be a useful tool to study dynamic alterations of the F-actin cortex in dependence of F-actin regulators, insulin or drugs that modulate insulin sensitivity.

To gain information about the internal properties of mature adipocytes, and in particular the lipid droplets themselves, we used a combined setup of ODT and Brillouin microscopy. When characterising the refractive index and Brillouin frequency shifts of different cellular compartments (lipid droplet, cytoplasm) of mature adipocytes, we observed a higher refractive index and Brillouin shift for lipid droplets compared to the cytoplasm, similar to our recent reports (29). A higher refractive index was also previously noted by Shoham et al. (30). However, due to the low density of the lipid droplet components, the higher refractive index was associated with lower longitudinal moduli, which shows clearly that the refractive index by itself does not serve as a proxy for the lipid droplets’ mechanical properties although this was recently suggested by Shoham et al. (30). When we compared refractive index, Brillouin shift and longitudinal moduli of lipid droplets between control and OA fed adipocytes, we did not find relevant differences between these different feeding conditions. This suggests that the mechanical differences between these feeding conditions revealed by AFM reflect differences in the lipid-to-cytoplasmic ratio and/or alterations in the cortical F-actin structures, which is supposedly the dominant part of the cytoskeleton probed by AFM indentation tests.

### Changes in cytoskeleton and tissue stiffness correlate with insulin resistance

In our study we found that tissues of obese animals were more compliant compared to lean controls, while increased tissue stiffness values were detected for diabetic animals. The finding of decreased tissue stiffness for obese (non-diabetic) tissues is in line with our *in vitro* findings of decreased apparent Young’s moduli for oleic acid fed cells with enlarged lipid droplets. It is, however, in contrast to a previous report by Mihai et al. Their study has compared the mechanical properties of epididymal fat tissue from obese (ob/ob) mice and wildtype mice C57BL/6 by shear rheometry to test the applicability of different mechanical models for adipose tissue. Although a comparison between lean and obese tissue was not the focus, the presented data suggest increased tissue stiffness for obese animals (10). The ob/ob mouse strain used in that study is obese due to excessive feeding and, depending on its age, hyperglycemic (the level of insulin resistance was not reported in that study though). In another study, tissue stiffness of mammary fat pads of ob/ob mice was found to be increased compared to lean control animals (11). However, experiments were conducted on rehydrated paraffin sections or cell-derived matrices, so the mechanical alterations at the cellular level were not directly addressed and the results are therefore not directly comparable to our study. Moreover, adipose tissues from different anatomic depots might be difficult to compare. Another significant factor differing among different studies may be animal age, since stiffness of several tissues has been demonstrated to increase with age (45). The lower apparent Young’s moduli detected by our AFM-indentation tests for (non-diabetic) obese compared to lean animals might reflect differences in the adipocytes’ cytoskeleton and/or ECM stiffness. Previous studies have, for instance, analysed changes in the F-actin cytoskeleton in obese mice, that had been on a high-fat diet for two weeks. Interestingly, increased F-actin levels were detected for the obese animals, from which one may at first expect increased stiffness values (46). In the mentioned study, however, the effects of high fat diet on tissue mechanics were not directly addressed. Also, changes in F-actin stainings and measured cortical thickness may not necessarily relate directly to changes in cortical stiffness, as recently reported for mitotic cells (47). Of course, we also have to consider that not only cellular components (e.g. other cell types found in adipose tissue), but also cell-cell contacts and importantly also extracellular matrix (ECM) contribute to tissue mechanics. This network of ECM fibrils surrounding adipocytes provides mechanical support, and, in particular fibrotic alterations typical to diabetes type II, are expected to increase tissue stiffness (48)(49)(50)(9). In line with that, recently a positive correlation between the extent of tissue fibrosis and stiffness was reported for human adipose tissue by Abdennour et al. (12). Thus, our finding of increased tissue stiffness for diabetic mice is in line with the study by Abdennour et al. (12) and also with a more recent report by Wenderott et al. (13). Moreover, Abdennour et al. probed subcutaneous white adipose tissue of human patients by vibration-controlled transient elastography. Adipose tissue fibrosis and diabetes were found to be associated with increased stiffness (12). It is feasible that the increased tissue stiffness of db/db mice we observed might be attributed to increased fibrosis, which might also explain the higher levels of local variations in stiffness values. To confirm this hypothesis, however, a more detailed characterisation of the ECM structures within the adipose tissue has to be performed, as well as functional tests systematically probing single components of the tissue.

It is well known that mechanical cues in the tissue can alter structure and function of tissue resident cells, and contribute to tissue development and homeostasis (51). In the context of adipose tissue, differentiation of preadipocytes has been shown to be modulated by mechanical cues provided by the cells’ microenvironment (15). Also, adipose tissue-derived stem cells cultured on substrates mimicking the stiffness of native adipose tissue (Young’s modulus, *E* = 2 kPa) increased their expression of adipogenic markers compared to stiffer substrates (17). In addition, while cells on compliant substrates differentiated more efficiently into adipocytes, they were more prone to differentiate towards the osteogenic lineage on stiff substrates, although this was dependent on soluble factors, suggesting a combinatorial regulation by both biochemical and mechanical cues (18)(19)(20). We observed the same stiffness-dependent behaviour of SGBS cells when culture on polyacrylamide (PAA) gels, where cells differentiated more efficiently towards adipocytes on hydrogels of lower elastic moduli (200Pa compared to 20kPa), adopted a more compliant phenotype and showed less nuclear YAP/TAZ translocation compared to stiff substrates (Supplementary Fig. 9). Within adipose tissues, preadipocytes and mature adipocytes are in contact with ECM molecules and neighbouring cells, which both provide mechanical cues to them. Thus, the seen differences in adipose tissue of lower stiffness for obese tissues might have an effect on the differentiation behaviour of neighbouring precursor cells, thereby balancing between hyperplasia and hypertrophy. Besides preadipocytes, also other tissue-resident cells including macrophages might be influenced by stiffness changes of their surroundings. In fact, it has been previously been shown that macrophages are mechanosensitive (52)(53)(54). Since chronic low-grade inflammation, mediated for instance by macrophages, has been associated with insulin resistance (55)(56), this might in addition contribute to the development of dysfunctional adipocytes in stiffened tissues.

### Conclusions and Outlook

Taken together, our results reveal a softening response of adipocytes not only at the onset of their differentiation but also during further maturation and lipid droplet size increase due to excessive lipid accumulation. These changes in mechanical phenotype with nutritional state were also reflected by a more compliant phenotype of adipose tissue after feeding animals a high fat diet, which was reversed, however, in diabetic animals. Altogether, our results provide new insights into the mechanical properties of adipose cells and tissue across different functional states, such as differentiation, nutritional state and diabetes type II. Based on these data, a mechano-sensory mechanism is proposed that could potentially regulate the balance between hyperplasia and hypertrophy to ensure healthy tissue expansion under nutrient excess. Our study sets the basis for more in-depths explorations of how changes in diabetic tissue mechanics and/or the resulting mechano-sensory responses could potentially be targeted to maintain tissue functionality.

## Methods

### Cell culture and adipogenic induction

Simpson-Golabi-Behmel Syndrome (SGBS) cells were originally prepared by Prof. Wabitsch’s group (University Medical Center Ulm, Germany) from an adipose tissue specimen of a diseased patient with SGBS (57, 58). A sample of subcutaneous adipose tissue (500 mg) was obtained post-mortem with informed consent from legal guardians. SGBS cells are a well-established human adipocyte *in vitro* model used in more than 200 labs world-wide. Cells were cultured as previously described (57). Cells were maintained in Dulbecco’s modified Eagles’ Medium (DMEM)/nutrient F-12 Ham supplemented with 4 μM panthotenic, 8 μM biotin (Pan/Bio), 100 U/ml Penicillin/100 μg/ml Streptomycin (= 0F Medium), together with 10% FBS (OF+FBS) at 37°C using T75 flasks. For adipogenic differentiation, cells were detached using TrypLE Express and seeded — depending on the experiment — onto glass bottom dishes (WPI, 35mm) (AFM), 6 well plates (western blots) or Thermanox coverslips (13 mm) (imaging) laid within 24-well plates and adipogenically induced as follows: After washing three times with serum-free DMEM/F12, cells were incubated for four days with OF Medium complemented with 10 μg/ml human transferrin (Sigma), 20 nM human insulin (Sigma), 2 μM Rosiglitazone, 100 nM dexamethasone (Sigma), 250 μM 3-isobutyl-1-1-methylxantine IBMX (Sigma), 100 nM Cortisol (Sigma), and 0.2 nM triiodothyronine T3 (Sigma). Then, the medium was exchanged to OF Medium supplemented with only transferrin, insulin, cortisol, T3 (concentrations as above). Medium was replaced every fourth day (58). To test the effect of drugs interfering with actomyosin contractility on differentiation, adipogenic medium was supplemented with 10 μM Y27632 or 400 nM Cytochalasin D (both from Sigma) from day 1 of adipogenic differentiation on. For oleic acid (OA) feeding experiments, differentiation medium was supplemented with either 60 μM OA (Sigma, BSA conjugated) or equivalent concentration of fatty acid free BSA from day 4 on. To test the effect of the ROCK inhibitor on lipid droplet size of mature adipocytes, 10 μM Y27632 was added in addition to OA from day 11 to 13.

### Animals

#### Fly culture

Wild type flies (*cantonS*) were cultured on standard food (Bloomington stock center, https://bdsc.indiana.edu) at 20°C maintaining a 12h day/night cycle. Eggs from flies kept on respective lipid defined diets were transferred onto fresh lipid defined food and cultivated at 20°C. Lipid defined food was composed as follows: 1. YAF-Oil: Yeast autolysate (Sigma Aldrich, 100g/l), Glucose (Roth, 100g/l), Agar-Agar (Roth, 10g/l), Nipagin (Sigma Aldrich, 1g/l) and cold-pressed olive oil (grocery store, 10g/l). 2. YAF-Sterol: Yeast autolysate (Sigma Aldrich, 110g/l), Glucose (Roth, 110g/l), Agar-Agar (Roth, 10g/l), Nipagin (Sigma Aldrich, 1g/l) and ß-Sitosterol (Sigma-Aldrich, 1g/l). Both, YAF-Oil and YAF-Sterol, allow Drosophila to complete multiple regenerative cycles at 20°C.

#### Mouse

Male C57BL/6-J mice (Janvier, Le Genest St Isle, France; 6 weeks of age) were housed three per cage at the Helmholtz Centre in Munich under standard controlled conditions. All procedures were in line with ARRIVE guidelines and approved by the Animal Use and Care Committee of Bavaria, Germany (ROB 55.2-1-54-2532-33-2014). To cause overweight and metabolic impairments, mice were ad libitum fed with high-fat diet (HFD, 58% calories from fat; 5.56□kcal/g, D12331, Research Diets, New Brunswick, NJ) or standard chow (Altromin, Germany, #1310, 14% calories from fat, 3.34 kcal/g) for 40 weeks. Ob/ob and db/db mice were from Janvier, and about 18 weeks old at the time of mechanical analysis. Mice were sacrificed by cervical dislocation (Landesdirektion Dresden AZ 24-9168.24-1/2013-17, DD24.1-5131/396/9). Gonadal adipose fat tissue was excised and immediately probed by AFM (native tissue) as described below or processed for histology. For Hematoxilin/Eosin or Masson’s Trichrome stainings, gonadal fat pads were washed in 1x PBS after excision and immediately fixed in 4% PFA buffered with PBS (Sigma). After 24h PFA was replaced with 1xPBS and samples were dehydrated and embedded in Paraffin.

### Atomic Force Microscopy (AFM) on single cells

For AFM indentation experiments on single SGBS and NIH3T3-L1 cells, a Nanowizard I or IV were used (JPK Instruments/Bruker Berlin). Arrow-TL1 cantilevers (Nanoworld) with a nominal spring constant of 0.035 – 0.050 N/m that had been modified with a polystyrene bead of 5 μm diameter (microparticles GmbH), were calibrated by the thermal noise method using built-in procedures of the AFM software. Cells growing on glass slides were gently rinsed with CO_2_-independent medium, which was also used for experiments. Then the cantilever was lowered with a speed of 5 μm/s onto the cells surface until a relative set point of 2.5 nN was reached and retracted. The resulting force distance curves were analysed using the JPK image processing software (JPK Instruments). Force-distance data were corrected for the tip-sample separation and fitted with the Hertz/Sneddon model fit for a spherical indenter to extract the apparent Young’s modulus (59, 60). A Poisson ratio of 0.5 was assumed. All experiments were performed at 37°C.

### Probing murine and fly adipose tissue stiffness by Atomic Force Microscopy (AFM)

Murine tissues were probed using a Cellhesion 200 equipped with a motorized stage (JPK Instruments/Bruker). Within 10 minutes after sacrificing, gonadal white adipose tissue (GWAT) was excised and transferred into CO_2_ independent medium (Life technologies). Tissue was trimmed and mounted onto glass-bottom Petri dishes (WPI), by attaching the edges to the glass surfaces using tissue seal (Braun). Tissue samples were carefully washed with CO_2_ independent medium (Life technologies), which was also used for experiments. Then a cantilever (Arrow TL1), which had been modified with a polystyrene bead (microparticles GmbH) of 10 μm diameter using epoxyglue (Faserverbundstoffe), was calibrated by the thermal noise method using built-in procedures of the AFM software. To perform indentation experiments, the cantilever was lowered at a speed of 10 μm/sec onto the tissue until a force of 2 nN was reached, and retracted again. For each animal usually both fat pads were probed. On each tissue section 8-10 areas of interest were probed using a 4 × 4 grid of 200 μm × 200 μm. Per condition in between 7 and 19 animals were analysed. Experiments were carried out at 37°C within 1.5 h after sacrificing the mouse.

For AFM indentation tests on fly fat bodies, feeding 3^rd^ instar larvae were collected, dissected in Graces medium at RT and fat bodies were transferred onto glass bottom dishes (WPI). Mechanical measurements started within 20min after tissue preparation. Tissues were probed using a Nanowizard 4 (JPK Instruments/Bruker) at room temperature (18-20°C) in Graces medium. During indentation experiments, the cantilever (same as above for murine tissue) was lowered at a speed of 5 μm/sec onto the tissue until a force of 4 nN was reached, and retracted again. For each animal both fat pads were probed if possible. On each tissue 5-8 regions of interest were probed using a 3 × 3 grid of 30 μm × 40 μm. Per condition in between 14 and 21 fat bodies were analysed. Force distance curves were processed as described above.

### Microrheology of adipose cells and tissues using AFM

AFM microrheology experiments were conducted on a Nanowizard 4 (JPK Instruments). PNP-TR-TL (Nanoworld) cantilevers with a nominal spring constant of 0.08 mN/m were modified with a polystyrene bead of 5 μm diameter for adipose cells and 10 μm diameter for adipose tissues (microparticles GmbH). The cantilever spring constants were measured prior each experiment using the thermal noise method implemented in the AFM software. For AFM microrheology experiments, the cantilever was lowered with a speed of 5 μm/s until a force set point of 2 nN for adipocytes and 4 nN for adipose tissues was reached. At an approximate indentation depth *δ*o of 1 μm, the cantilever was then oscillated by a sinusoidal motion of the piezo elements at an amplitude of 10 (or 30) nm for a period of 10 cycles. The procedure was repeated for different oscillation frequencies for each cell in the range of 3 – 200 Hz for adipocytes and 5 – 60 Hz for adipose tissues. In each oscillatory measurement the whole frequency range was applied at the same spot and three repetition were done for each cell in different regions. For post-processing, the force and indentation signals were fitted using a sinusoidal function to extract the amplitude and phase angle of each signal. Data were analyzed analogously to the procedure described by Alcaraz et al. (61) but for a spherical not a pyramidal indenter. Briefly, the method relies on the linearization of the Hertz model for a spherical indenter due to small oscillations by using the first term of the Taylor expansion and subsequent transformation to the frequency domain:

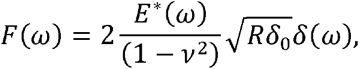

where *F*(*ω*) and δ (*ω*) are the force and indentation signals in the frequency domain, respectively, *E*\*(*ω*) is the complex Young’s modulus, *v* is the Poisson’s ratio assumed to be 0.5, *R* is the radius of the indenter and *ω* is the angular frequency. The complex shear modulus *G*\*(*ω*) can be written using 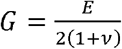 (62):

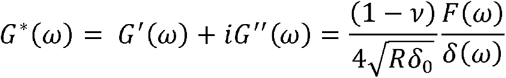

where *G ′*(*ω*) is the storage modulus and *G ″*(*ω*) is the loss modulus. The ratio of the force *F*(*ω*) and indentation *δ*(*ω*) is calculated from the measured amplitudes *A*^*F*^(*ω*) and *A*^*δ*^ (*ω*) and the phase shifts *θ* ^*F*^(*ω*) and *θ*^*δ*^ (*ω*) of the oscillatory signals (63):

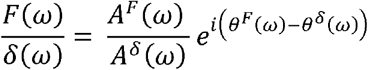

where the difference of the phase shifts (*θ* ^*F*^(*ω*)– *θ*^*δ*^ (*ω*)) is in the range of 0° (elastic solid) and 90° (viscous fluid). Furthermore, the hydrodynamic drag contribution on the cantilever oscillation was estimated and subtracted from the complex shear modulus as previously described (63)(64):

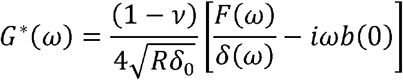

Where *b*(*h*) is the hydrodynamic drag coefficient function measured from non-contact oscillations of the cantilever at different distances *h* from the sample, and *b*(0) is the extrapolation to distance 0 from the sample. For PNP-TR-TL cantilevers, the hydrodynamic drag coefficient was estimated to be 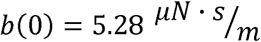.

### Cell staining and image analysis

For analysis of cell morphology or percentages of cells with lipid droplets, cells were fixed at the indicated timepoints using 4% formaldehyde solution in PBS (pH 7.4) for 5-10 minutes, washed with PBS and stained with 0.4 μM Phalloidin-TRITC (Sigma) or Phalloidin-Alexa 647 (Cell signalling), 0.1μg/ml nile red (Sigma), and 5 μg/ml DAPI in blocking buffer (PBS, 2% BSA). For YAP/TAZ staining, cells were permeabilised after fixation using 0.2%TritonX/PBS and blocked with 2% BSA/PBS for 15 minutes. Then, they were incubated with 5 μg/ml primary monoclonal antibody to YAP/TAZ (Cell signalling, clone D24E4) for 60min, followed by a washing step in PBS and 60min incubation with C5-conjugated secondary antibodies (Jackson Immunoresearch). Samples were mounted using mounting medium (Zeiss) and viewed using a confocal fluorescence microscope (Zeiss LSM 700 or 780) using a 40x water immersion objective (Zeiss). Stress fibers were quantified in confocal images of F-actin stained SGBS cells using Fiji(65). A cross-section was taken (perpendicular to the stress fibres orientation) and maxima were counted. F-actin fibril anisotropy was measured using the Fibril Tool plugin for the NIH ImageJ program, as described by Boudaoud et al (66). Cells and the image channel of F-actin were manually selected. Lipid droplet sizes were analysed in confocal fluorescence images of nile red stained cells using Ilastik version 1.3 (pixel and object classification) (67). YAP/TAZ localisation was determined based on intensity levels in confocal fluorescence images of YAP/TAZ, phalloidin-TRITC and DAPI stained cells using FIJI.

### Western blot analysis

Cells were lysed using Laemmli sample buffer (62.5 mM Tris-HCl (pH 6.8), 2% SDS, 10% glycerol, 0.01% bromophenol blue). Samples were boiled (5 min 95 °C), loaded on gradient gels (MiniProtean, Biorad), and separated by reducing SDS-PAGE. After blotting onto a PVDF membrane (Merck Millipore), membranes were blocked with TBS-Tween (20 mM Tris, 137 mM NaCl, 0.1% Tween) containing 5% (w/v) non-fat dry milk for 1 hour. Membranes were incubated overnight at 4°C (5% BSA/PBS) with antibodies against pMLC (ser19), ppMLC (Thr18/ser19), PPARgamma antibodies (all three from Cell Signalling), tubulin (Abcam), GAPDH (Abcam). After incubation with respective HRP-conjugated secondary antibodies (Abcam), chemiluminescence detection was performed using Enhanced Chemi-Luminescence (ECL) (Thermofisher). Densitometric analysis was done using Fiji.

### MRE on murine adipose tissue

A 0.5 Tesla permanent magnet MRI scanner (Pure Devices GmbH) was used for the MRE studies. The scanner controlled a custom piezoelectric driver (Piezosystem Jena) mounted on a glass tube with an inner diameter of 8.0 mm and a length of 15 cm, the lower end of which was placed in the magnet(68). The cylindrical tissue samples with a diameter of 8 mm and a height of about 3 to 5 mm were placed at the bottom of the glass tube. The tube was sealed with a PVC plug and a PVC pad, where the pad ensured shear wave excitation from the cylinder walls into the tissue. Five drive frequencies of 400, 600, 800, 1000 and 1400 Hz were applied and encoded by a motion-sensitized phase-contrast imaging sequence as detailed in Braun et al (69). In brief, the following acquisition parameters were used: repetition time = 2000 ms, echo time = 42 ms, slice thickness = 3 mm, matrix size = 64 × 64, field of view = (9.6 × 9.6) mm^2^, voxel size = (0.15 × 0.15 × 3) mm^3^. The acquired phase data were unwrapped and Fourier-transformed in time to extract complex-valued wave images at driving frequency *f*. All data points of the *z*-deflection (parallel to the cylinder axis) were mapped onto cylindrical coordinates and averaged over the azimuthal angle, yielding a single profile along the radial coordinate. The resulting profiles were fitted by the analytical solution of shear waves in a z-infinite cylinder as described previously where the Bessel function of first kind yields the desired result of the fit, the complex wave number *k*\* = *k*’ + i*k*’’(69). As described by Tzschätzsch et al.(70), we translated k* into two real-valued quantities related to elastic stiffness (shear wave speed *c*) and inverse attenuation (shear wave penetration rate *a*), both in the dimension of m/s:

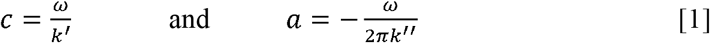

where *ω* is the angular driving frequency equal to 2π*f*.

As described in a previous study we directly fitted *c* and *a* by a viscoelastic model to derive shear modulus-related parameters instead of first calculating a complex shear modulus *G*\* from *k*\* and then fitting *G*\*(71). We chose this approach because *c* and *a* are the primary results of the Bessel fit instead of *G*\*, which is normally obtained from direct inversion–based MRE.

*c* and *a* were converted to *G\** by the following equation:

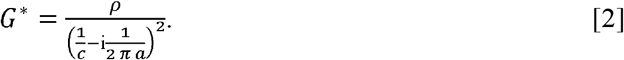

### Fluorescence/ODT/Brillouin microscopy

Measurements of the fluorescence intensity, refractive index (RI) and Brillouin shift were done using a custom build optical setup that combines epi-fluorescence imaging, optical diffraction tomography and Brillouin microscopy in a single device (29). In order to acquire the RI, the sample is illuminated under multiple different angles by a collimated beam from a laser source with a wavelength of 532 nm. The transmitted beam interferes on a camera chip with a reference beam. The resulting spatially modulated holograms are evaluated to reconstruct the three-dimensional refractive index distribution of the sample. For Brillouin microscopy, the sample is illuminated with a focused beam. The backscattered light is collected by a single-mode fibre which acts as pinhole confocal to the illumination optics. The fibre guides the light towards a two-stage VIPA spectrometer to measure the Brillouin frequency shift (72). The setup achieves a spatial resolution of 0.25 μm in the lateral plane and 0.5 μm in the axial direction for ODT and 0.4 μm in the lateral plane and 1 μm in the axial direction for Brillouin microscopy.

## Statistical analysis

GraphPad Prism was used to plot data, with the exception of violin plots that were generated using the R package Vioplot (73). Statistical tests were performed in Graphpad Prism. A two-tailed significance level of 5% was considered statistically significant (p<0.05). The respective tests applied are indicated in figure legends.

## Acknowledgements

We thank all Guck lab members, in particular Elke Ulbricht, Stephanie Möllmert, Dominik Eberle, Kaddour Bounab and Isabel Richter for scientific discussions and help with experiments. We also thank JPK Instruments/Bruker, in particular Jörg Barner, for technical support. We also thank Prof. Aránzazu del Campo and her team for providing materials for production of PAA gels. This project has received funding from the Alexander-von-Humboldt Stiftung (Humboldt-Professorship to J.G.), the Marie-Curie ITN BIOPOL (J.G.), the Deutsche Forschungsgemeinschaft DFG (419138906 to J.G., Project-ID 329628492 – SFB 1321 to K.S., and BR54890/2-1 to M.B.), and the Volkswagen Stiftung (grant agreement number 92847 to J.G.). A.T. is a fellow of the MSNZ supported by the Deutsche Krebshilfe. We are grateful to the MPI-CBG, CRTD and Helmholtz mouse house staff for support. We thank the light microscopy facility of the CMCB (in part funded by the State of Saxony and the European Fund for Regional Development-EFRE).

## Contributions

The project was conceived by K.S., P.K., J.G. and A.T. Cell and tissue mechanical experiments by AFM, Brillouin, ODT and MRE were performed by S.A., R.S., A.S., P.K., A.H., K.K. and A.T. AFM and Brillouin data analysis was performed by S.A., R.S. and A.T. Mouse tissue preparation and characterisation was done by P.K. and A.H. Fly manipulation and tissue preparation was done by M.B. SGBS cells were provided by M.W. ODT and Brillouin microscopy setups were built by R.S. and K.K. MRE setup and data analysis were done by A.S, J.B., and I.S. PAA gels were prepared by J.C.E. The draft of the manuscript was written by A.T. with subsequent contributions from all co-authors, in particular S.A., P.K., J.G. and K.S.

