## Supplementary Information for "Adipose cells and tissues soften with lipid accumulation while in diabetes adipose tissue stiffens"

#### Supplementary figures

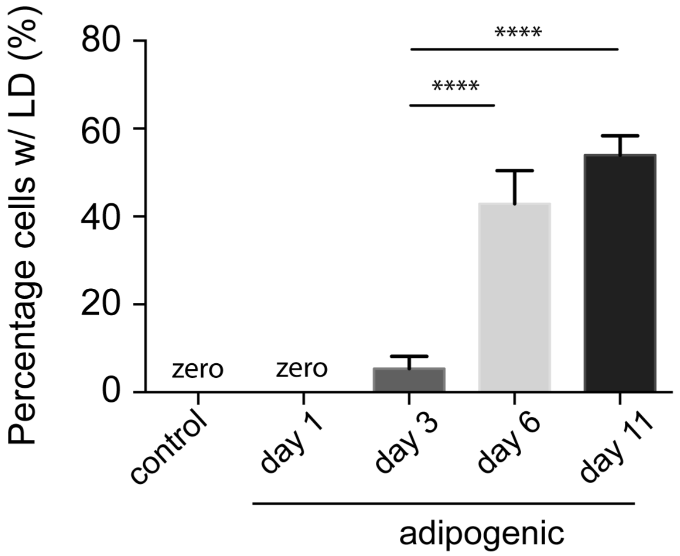


**Supplementary Figure 1.** Representative experiment with differentiating SGBS cells on day 1, 3, 6 and 11 after adipogenic induction. Cells were stained for F-actin (Phalloidin-TRITC, red), Lipid droplets (nile red, green), and nuclei (DAPI, blue) (as in Figure 1), and the percentage of cells with lipid droplet accumulation was determined in 6 images per condition. For statistical analysis, a one-way ANOVA with a Tukey's multiple comparisons test were performed. **** indicates p<0.0001.

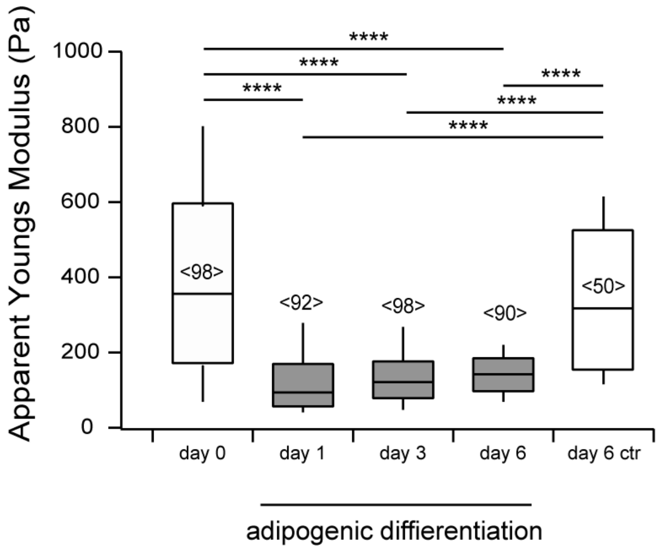


**Supplementary Figure 2.** Mechanical characterisation of NIH3T3-L1 cells by AFM indentation experiments using a spherical indenter (5 µm diameter). Apparent Young's modulus data are presented as box whisker plots. For comparison a Kruskal-Wallis test with a Dunn's multiple comparisons test was performed. **** denotes p-values <0.0001.

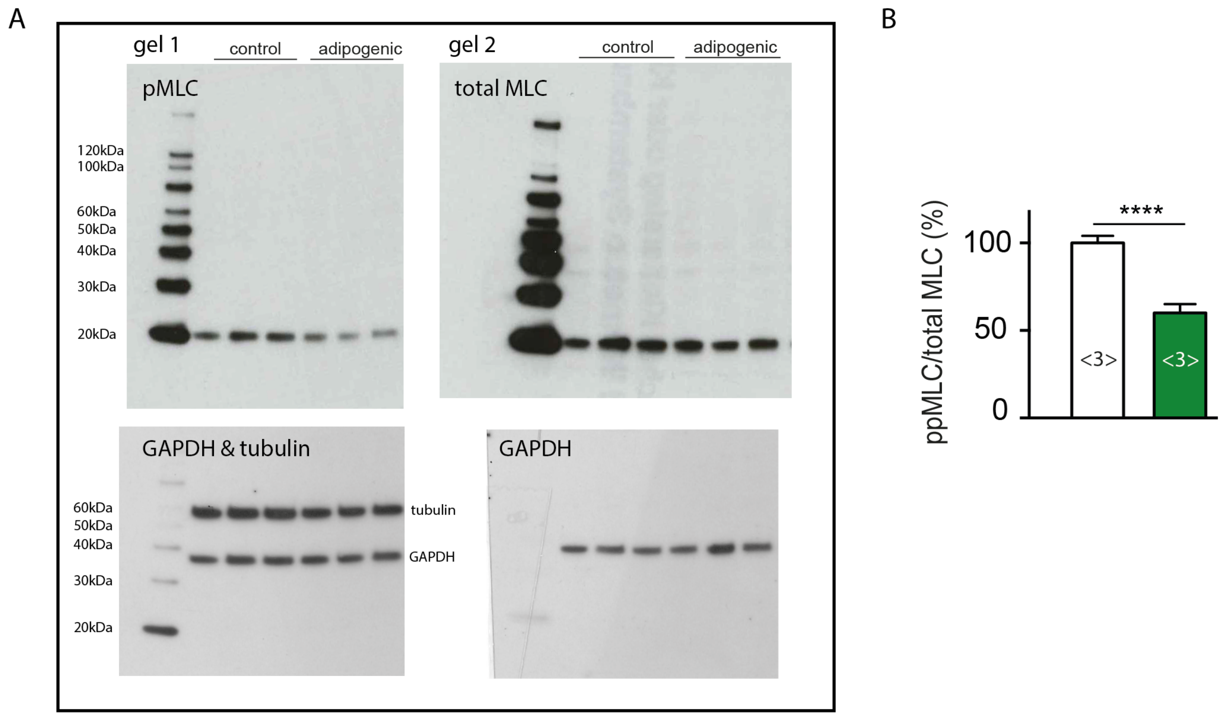


**Supplementary Figure 3. (A)** Full-length western blot corresponding to Fig. 3F, of SGBS lysates probed for monophosphorylated (pMLC2, Ser19, #3671, Cell Signaling) and total myosin light chain 2 (MLC2) levels (D18E2, cell signalling) at day 1 (control or adipogenic medium). As loading control GAPDH levels were quantified. (**B**) Western blot analysis of SGBS lysates for di-phosphorylated (ppMLC2, Thr18/Ser19, #3674, Cell Signaling).

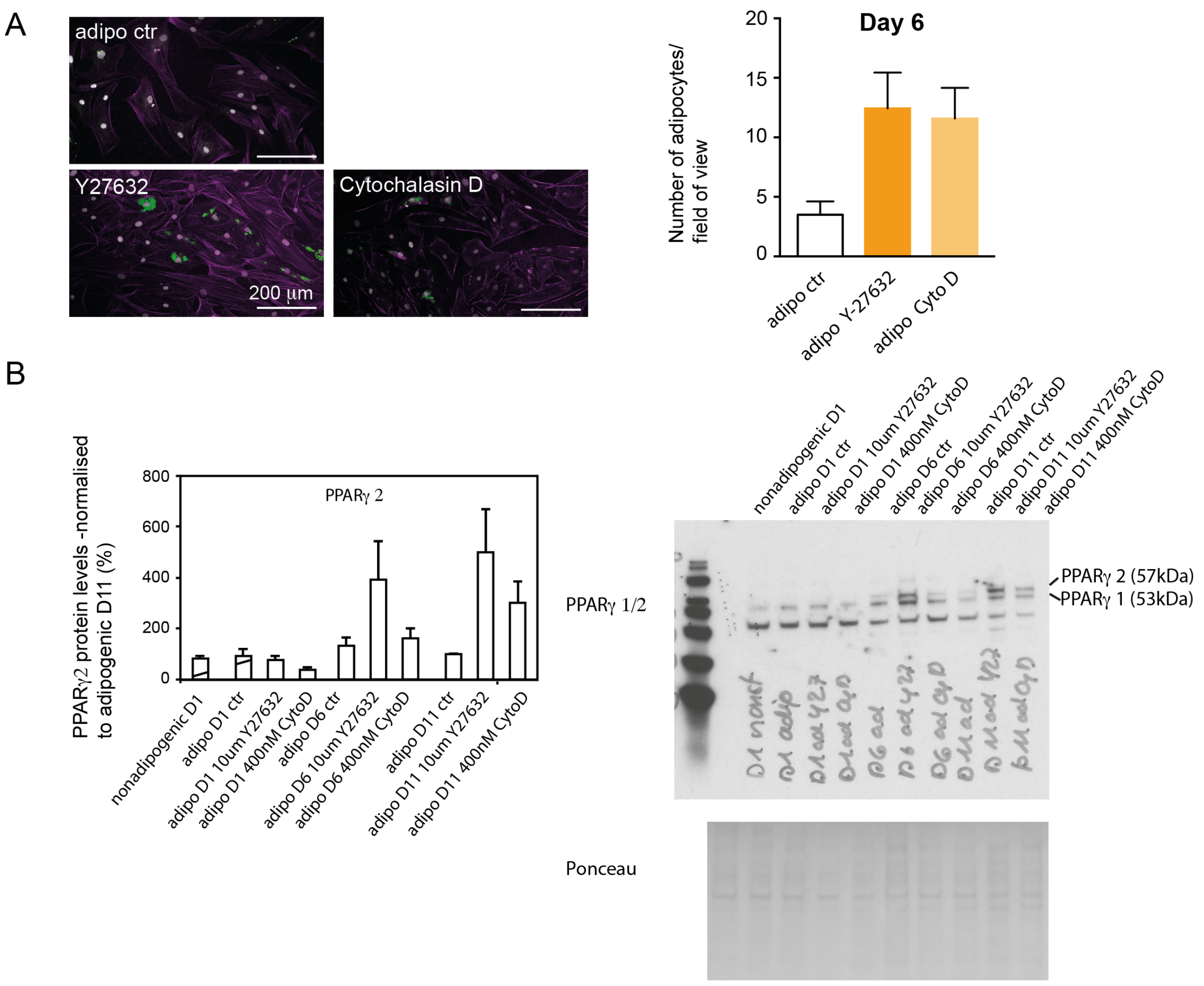


**Supplementary Figure 4. (A)** Representative confocal microscopy images of SGBS cells (day 6 adipogenic) stained for F-actin (Phalloidin-Alexa647, purple), lipid droplets (nile red, green), and nuclei (DAPI, white). Right: Quantification of the number of cells with lipid droplets per image for adipogenic controls (vehicle control), and cells treated with 10 µM Y-27632 or 400 nM Cytochalasin D (Cyto D). Mean +/- Standard error on the mean is shown. **(B)** Western blot analysis of SGBS lysates for PPARγ after drug treatment (10 µM Y-27632, 400 nM Cytochalasin D). Exemplary western blot on the right.

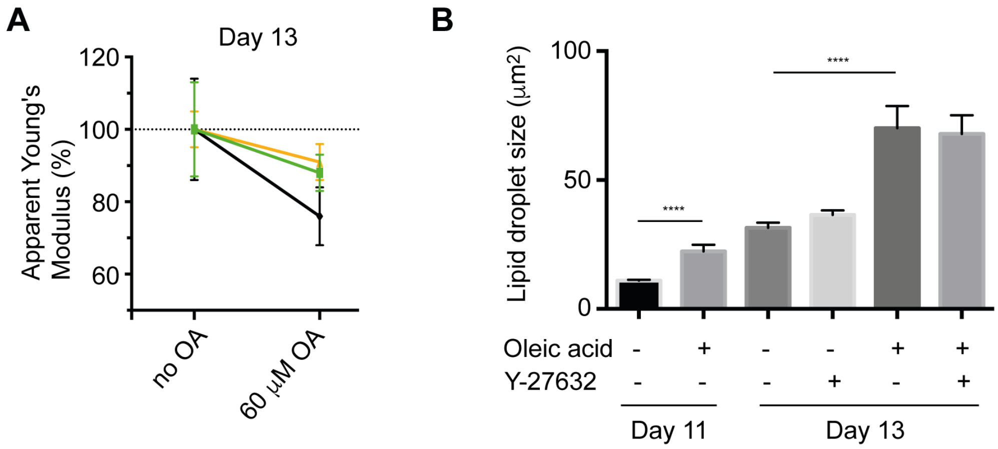


**Supplementary Figure 5. (A)** Three independent AFM indentation experiments on SGBS cells after oleic acid feeding on day 13 using a spherical indenter (5 µm diameter). Apparent Young's modulus data are presented (mean +/- standard error on the mean). **(B)** Lipid droplet size after oleic acid feeding in presence and absence of 10 µM Y-27632. Data are presented as mean +/- SEM.

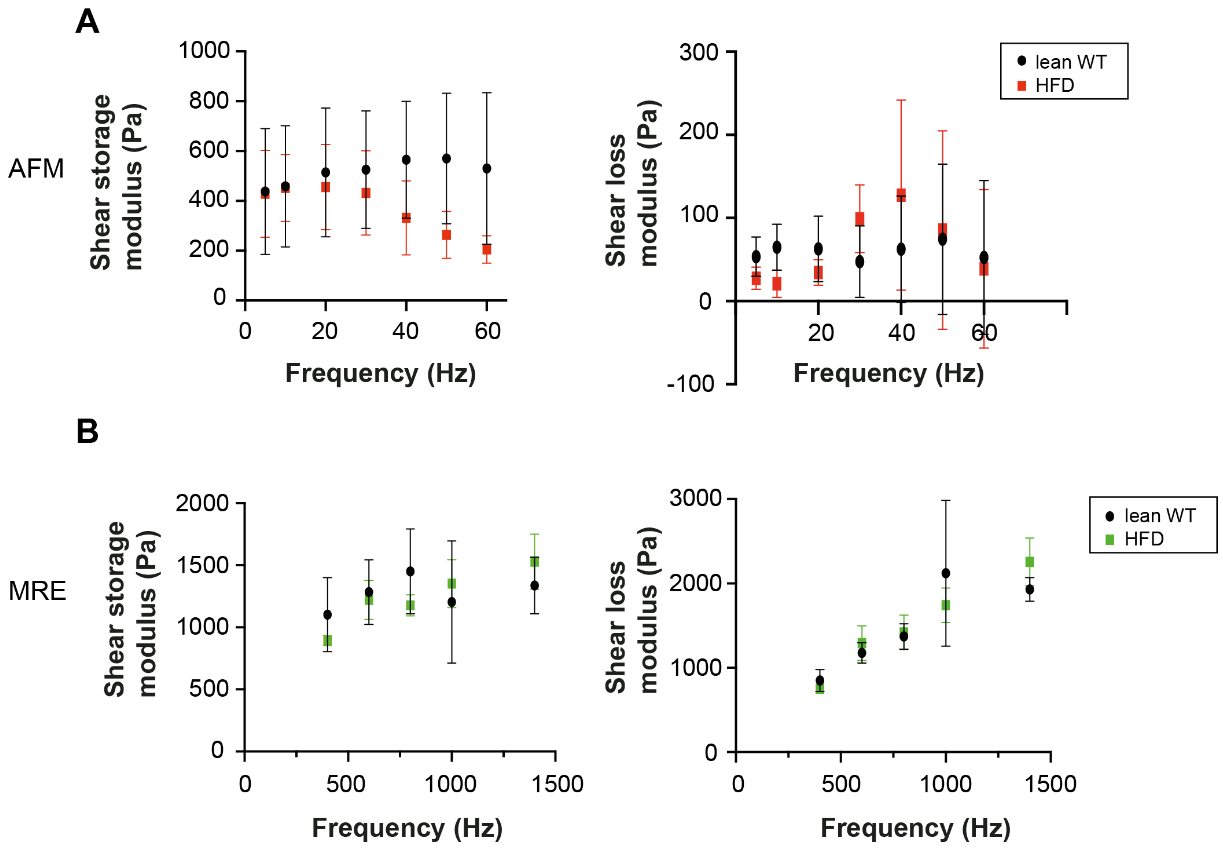


**Supplementary Figure 6. Dynamic measurements on gonadal mouse adipose tissue (A)** AFM microrheology data over a frequency range of 5 to 60 Hz. Shear storage (G') and loss moduli (G'') are presented as mean+/- SEM. Adipose tissue was probed locally using a spherical indenter (5 µm diameter) for 4 lean and 4 high fat diet fed animals **(B)** Shear storage and loss calculated from MRE data over a frequency range of 400 to 1400Hz. 6 animals were analysed per condition.

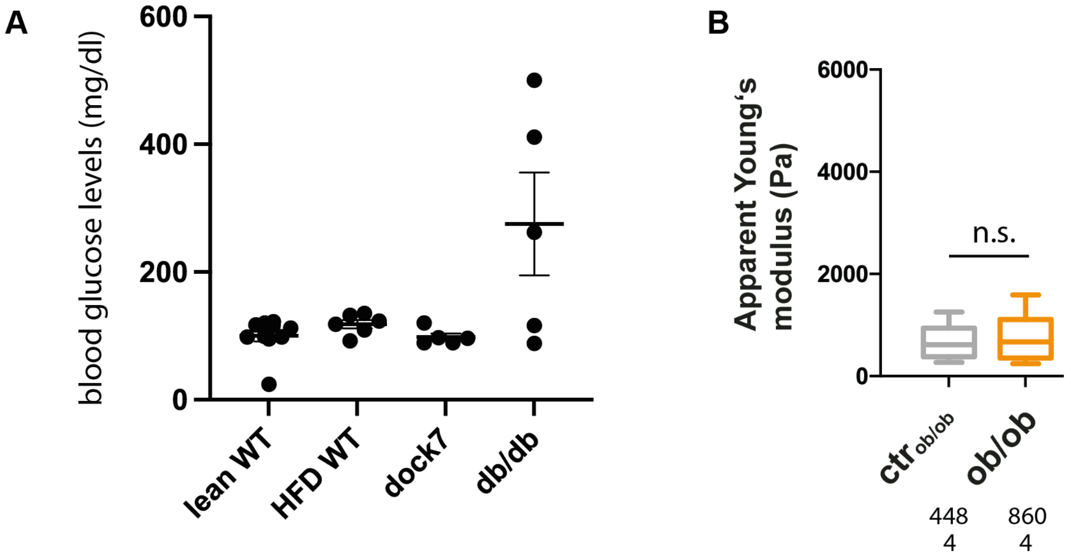


**Supplementary Figure 7. (A)** Blood glucose levels measured from chow and high fat diet (HFD) fed wildtype (lean and HFD WT), and dock7 and db/db mice on normal diet. **(B)** AFM indentation experiment on gonadal mouse adipose tissue of leptin- deficient ob/ob mice. Apparent Young's modulus data are presented as box whisker plots. Number of analysed force-distance curves (above) and mice (below) are indicated below. Data were compared to respective controls by a Mann-Whitney test. **** indicates p-values <0.0001.

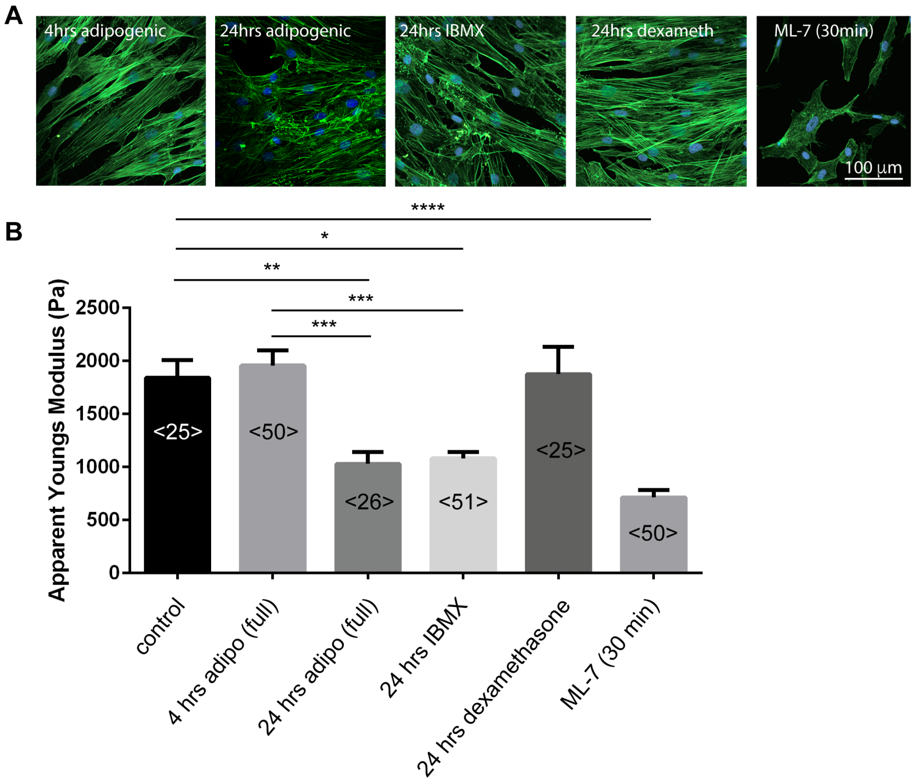


**Supplementary Figure 8.** Mechanical characterisation of SGBS cells by AFM indentation experiments using a spherical indenter (5 μm diameter). Cells were treated with full adipogenic medium, single components of it (250 μM IBMX, 100 nM Dexamethasone) or the myosin light chain kinase inhibitor ML-7 (10 μM). Apparent Young's modulus data are presented as bar plots. For comparison a Kruskal-Wallis test with a Dunn's multiple comparisons test was performed. * denotes p-values <0.05, ** denotes p-values <0.01, *** denotes p-values <0.001, **** denotes p-values <0.0001.

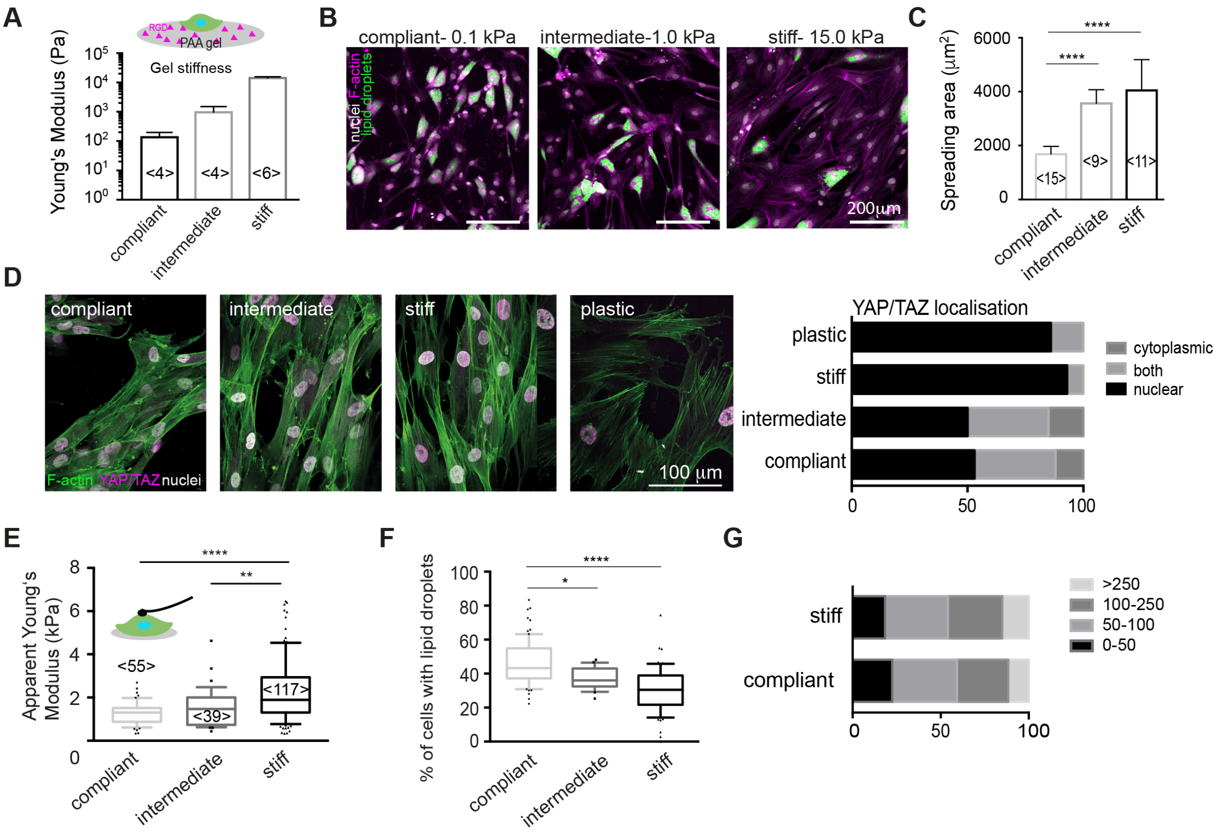


**Supplementary Figure 9.** SGBS cultures on elastic polyacrylamide (PAA) gels. (A) Mechanical characterisation of PAA gels by by AFM indentation experiments using a spherical indenter (5 µm diameter). Young's moduli were derived from force distance curves and plotted as bar charts (mean +/- SEM). (B) Representative confocal images of SGBS cells on PAA gels of different stiffness, stained for F-actin, nuclei and lipid droplets. (C) Quantification of cell spreading area using FIJI. (D) Confocal images of SGBS cells stained for YAP/TAY (antibody), F-actin (Phalloidin TRITC) and nuclei (DAPI) on different stiff PAA gels. Right: quantification of YAP/TAZ localisation. (E) Mechanical characterisation of SGBS cells grown on gels for 24hrs by AFM indentation using a spherical indenter (5 µm diameter). (F) Quantification of cells with lipid droplets on PAA gels of different stiffness. (G) Lipid droplet size distribution (in µm²) of SGBS cells on PAA gels on day 11 after adipogenic differentiation. For comparison a Kruskal-Wallis test with a Dunn's multiple comparisons test was performed. * denotes p-values <0.05, ** denotes p-values <0.01, *** denotes p-values <0.001, **** denotes p-values <0.0001.

### Supplementary tables

| | % acrylamide (w/v) | % bis-acrylamide (w/v) | % TEMED (v/v) |
|---|---|---|---|
| Soft | 5 | 0.07 | 0.3 |
| Intermediate | 7.5 | 0.06 | 0.3 |
| Stiff | 12 | 0.2 | 0.3 |

Supplementary Table 1: Composition of polyacrylamide hydrogel premixes

### Supplementary Methods

#### Preparation of polyacrylamide (PAA) gels

To prepare PAA gels of comparable ligand density at different stiffnesses, we used a previously described method (73), but additionally incorporating methylsulfonyl groups into the PAA hydrogel structure for later functionalisation with RGD ligands (74). Briefly, glass coverslips (13mm ø, Marienfeld) were first activated as follows: firstly, they were washed with 1N NaOH for 30 minutes, then with dd$H_2O$, ethanol and dd$H_2O$, and dried. Thereafter, they were amino-silanised using a solution of chloroform with 0.1% (v/v) triethylamine and 0.1% (v/v) allyltrichlorosilane (all from Sigma) for 30 minutes. Glass slides were again washed with dd$H_2O$ and incubated for 30 minutes with 0.5 % of glutaraldehyde solution (in dd$H_2O$). After a final wash with ddH2O, glass slides were dried and used within 2 days for the preparation of gel layers. To prepare polyacrylamide hydrogels, acrylamide, bis-acrylamide, PBS and TEMED were mixed according to Supplementary table 1 and de-gassed for 30 minutes under vacuum in a desiccator. Then, to 100 µl of that mix, 1 µl of 10% APS (dd$H_2O$) was added and mixed with a pipette. 80 µl of that mix were then combined with 10 µl of a 32 mg/ml solution of methylsulfone (in N,N-Dimethylformamide, DMF) and mixed again. Quickly, before polymerisation, droplets of 9.3 µl were added onto an ethanol-cleaned foil and sandwiched with the above-described activated glass slides. After 30 minutes at RT, hydrogels attached to the cover glasses were carefully detached from the foil, washed and kept in dd$H_2O$ until following RGD functionalisation. Then, 250 µl of a 0.5 mg/ml RGD/dd$H_2O$ (Pepnet) solution were added onto the gel surfaces and incubated overnight at RT. Afterwards, surfaces were washed 3 times with PBS, followed by a washing step with medium. Finally, cells were seeded as described above.

**Oil red staining**

Adipocytes were differentiated within 24 wells for 11 days and fixed for 30 min with 4% formaldehyde/PBS. To prepare the fresh oil red staining solution, 3 parts of a stock solution (250 mg oil red (Sigma) in 50 ml isopropanol) were mixed with two parts $ddH_20$. After letting it sit for 10 min, the solution was passed through a sterile filter (0.45 µm, Millex). Thereafter, 1 ml 60 % isopropanol was added to cell layers for 5 minutes. Isopropanol was then replaced by 0.5 ml oil red staining solution, gently added dropwise into the well. After 1 hour incubation at RT on a rocking shaker, cell layers were gently washed three times with $H_2O$ and dried. Cells were viewed using a stereomicroscope (Olympus) and images were taken. For quantitative oil red staining, oil red dye was extracted using 250 µl of 100% isopropanol, and 100 µl were transferred into a 96-well plate and absorbance was read on a plate reader at 520 nm (Tecan plate reader).
